# A palette of fluorophores that are differentially accumulated by wild-type and mutant strains of *Escherichia coli*: surrogate ligands for bacterial membrane transporters

**DOI:** 10.1101/2020.06.15.152629

**Authors:** Jesus Enrique Salcedo-Sora, Srijan Jindal, Steve O’Hagan, Douglas B. Kell

## Abstract

Our previous work had demonstrated that two commonly used fluorescent dyes that were accumulated by wild-type *E. coli* MG1655 were accumulated differentially in single-gene knockout strains, and also that they might be used as surrogates in flow cytometric transporter assays. We summarise the desirable properties of such stains, and here survey 143 candidate dyes. We triage them eventually (on the basis of signal, accumulation levels, and cost) to a palette of 39 commercially available and affordable fluorophores that are accumulated significantly by wild-type cells of the ‘Keio’ strain BW25113, as measured flow cytometrically. Cheminformatic analyses indicate both their similarities and their (much more considerable) structural differences. We describe the effects of pH and of the efflux pump inhibitor chlorpromazine on the accumulation. Even the ‘wild-type’ MG1655 and BW25113 strains can differ significantly in their ability to take up such dyes. We illustrate the highly differential uptake of our dyes into strains with particular lesions in, or overexpressed levels of, three particular transporters or transporter components (*yhjV, yihN*, and *tolC*). The relatively small collection of dyes described offers a rapid, inexpensive, convenient and valuable approach to the assessment of microbial physiology and transporter function.

## Introduction

Notwithstanding that the entire genome of strain MG1655 of *Escherichia coli* K12 was sequenced more than 20 years ago [1], some 35% of its genes are still of unknown function [2]. These genes are known as y-genes [3], and transporters are pre-eminent among them [2]. An important general problem [4, 5] thus involves the related questions of (i) what are the potential substrates for a given transporter, and (ii) what are the membrane transporters for an molecule that cells take up and/or efflux? Our focus here is on finding methods to help provide answers to these questions, in particular for the (approximately) 124 y-genes of *E. coli* that are considered from sequence analysis to be transporters.

Fluorescence flow cytometry provides a convenient and high-throughput means for assessing the extent of uptake and accumulation of a given fluorophore (e.g. [6-16]), and in recent work [17, 18] we have found this to be true in *E. coli in vivo*. In particular, the availability of the ‘Keio’ collection of single-gene knockouts in ‘non-essential’ genes [19, 20] allowed up to assess the contribution of many of these genes to uptake. The chief findings were that the carbocyanine dyes diSC3(5) and SYBR Green I could be taken up and/or effluxed differentially relative to the reference strain by a very great number of strains harbouring single-gene knockouts, consistent with the views that (i) any electron-transport-mediated membrane potential was not largely responsible for their steady-state uptake [21-23], and (ii) any non-transporter-mediated transport through the phospholipid bilayer was negligible [24-29].

The fluorophores chosen in this earlier work [18] were two dyes that are widely used in the assessment of the numbers and physiological status of intact bacteria, as they seem to permeate wild-type membranes more or less easily (presumably via a variety of transporters). While the strong dependence on transporter activity meant that such dyes did not faithfully reflect either bioenergetic parameters or nucleic acid content, we noted [18] that they opened up the possibility of high-throughput screening of transporter activity, including of competitive or inhibitory (non-fluorescent) substrates of membrane transporters (as in [30] for uptake and [31-36] for efflux).

The desirable properties of dyes (and assays) of this type include the following:

1. They are taken up intracellularly more or less rapidly by the target species
2. They do not interfere significantly with the host’s biochemistry at the concentrations used and on the timescale of interest
3. They have a high fluorescence signal (usually involving a high absorbance at the excitation wavelength and a high quantum yield)
4. They are not interfered with by autofluorescence (often implying a large Stokes shift or an absorbance nearer the red)
5. They show a fluorescence that is linear with intracellular concentration at the time point(s) of interest (consistent with point 2)
6. Assays should be done at constant cell number or (if they are not) that the number of cells used does not materially affect the external concentration of the dye
7. When used as a surrogate for a particular transporter, the bulk of the flux of interest is mediated by that transporter

We also recognise that in some cases (not least when binding is to DNA or to metal ions) the binding of such fluorophores to intracellular targets can induce changes in both the magnitude and the fluorescence spectrum of a given dye; this will in some cases need to be considered when the uptake of these dyes is used as a surrogate measure for transporter activity. It goes without saying that the expression of particular transporters (or anything else) depends on the growth conditions used, and these are typically controlled in *E. coli* by transcriptional regulatory networks (e.g. [37-40]). Consequently in any given condition there is a danger of ‘false negatives’ [41, 42] i.e. rejecting dyes that under other circumstances might be taken up well. Thus, we have sought to be inclusive of potentially useful dyes so far as is possible

Dyes with most or all of the desirable properties given above are not particularly common as applied to unfixed cells, and we recognised that a wider survey of potentially useful, bacterial-membrane-permeating dyes of different structures might prove of considerable value. Further, most dyes applied to biological cells are used on mammalian cells, and so may not be pertinent to bacteria. Nonetheless, we surveyed the catalogues of a wide variety of biological stain suppliers, plus other sources such as food colours, laser dyes, dyes used in the water industry for tracking, and even the core scaffolds used in organic light-emitting diodes. Thus, the present study represents an initial survey to this end, and provides a significant number of dyes that do seem to have most or all of the desired properties in the reference strain. In addition, we assess their utility in profiling membrane transporters using appropriate mutants.

## Methods

*E. coli* strain **BW25113** (Keio collection reference strain: *Δ(araD-araB)567, ΔlacZ4787*(::rrnB-3), *λ*^*-*^, *rph-1, Δ(rhaD-rhaB)568, hsdR514*) and certain other representatives of the Keio collection [19, 20] used here included the following gene knockouts: ***yhjV*** (*F-, Δ(araD-araB)567, ΔlacZ4787(::rrnB-3), λ-, ΔyhjV722::kan, rph-1, Δ(rhaD-rhaB)568, hsdR514*); ***yihN*** (*F-, Δ(araD-araB)567, ΔlacZ4787(::rrnB-3), λ-, rph-1, ΔyihN736::kan, Δ(rhaD-rhaB)568, hsdR514*); and ***tolC*** (*F-, Δ(araD-araB)567, ΔlacZ4787(::rrnB-3), λ-, ΔtolC732::kan, rph-1, Δ(rhaD-rhaB)568, hsdR514*). We also used the strain overexpressing *yhjV* from the ASKA collection: *E. coli* K-12, strain AG1 [*recA1 endA1 gyrA96 thi-1 hsdR17 (r K− m K+) supE44 relA1*] carrying recombinant constructs in the IPTG inducible plasmid pCA24N (CmR, *lacIq*). The induction with IPTG was optimised to 250µM for 3 hours (37°C, shaking 200 rpm) previous to the fluorophore uptake assays with the ASKA strain. Both the Keio and the ASKA collection [pmid16769691] were provided by the National Institute of Genetics, Mishima, Shizuoka, Japan).

For each fluorophore uptake assay the starting point was the spread of a small flake from a frozen culture stock of *E. coli* onto a fresh plate of complex solid media (Merck LB 110283) containing selective antibiotics when appropriate. Kanamycin was used at 50 µg/ml final concentration for the Keio knockout strains and chloramphenicol at 30 µg/ml for the ASKA strain). A single colony was then incubated in liquid complex media (Merck LB 110285) overnight cultures, 37°C, shaking 200 rpm in the absence of antibiotics. The overnight cultures were diluted 1:5000 in fresh liquid complex media and grown for 2 hours at 37°C shaking at 200 rpm. The cell density was then adjusted to ∼2000 cells.µL^-1^ as judged by turbidity (OD600). Cells were then exposed to fluorophores (37°C, 15 min or as indicated, with shaking at 1300 rpm) in 384-well plates and final volumes of 50µL.

The experiments with chlorpromazine (CPZ) were performed using two different protocols. For the first set of experiments, the protocol was replicated from [18], where 5µL CPZ (1mM in DMSO) were added to the wells of a 96-well plate in triplicate. A vacuum centrifuge was used to dry all the DMSO in the plates. Thereafter, 200µL of overnight-cultured *E. coli* (MG1655 or BW25113) were added to each well at 1000 cells.µL^-1^ final concentration in complex media. The plates were sealed and incubated at 30°C with 900 rpm shaking for 30 minutes. Then, DiSC3(5) or SYBR Green I were added to final concentration of 3µM and 1X (a 10,000-fold dilution of the stock material supplied), respectively [18]. The plate with DiSC3(5) was incubated for 2 minutes while the one with SYBR Green I was incubated for 15 minutes at 37°C before sampling in the flow cytometer. For the second set of experiments, CPZ was added to overnight-cultured *E. coli* (BW25113; 2000 cells.µL^-1^) cells at a final concentration of 10µM in complex media. This culture was then incubated with different fluorophores (3µM) at different pH values for 15 minutes. To modify the pH of the media the following buffers were added individually (cf. [43]) to the media at final concentrations of 100mM: MES (pH 6, 6.5), MOPS (pH 7.0, 7.5, 8.0) and Bicine (pH 8.5). Fluorophores were purchased from Sigma Aldrich, ThermoFisher, TCI America, or ATTO-TEC GmbH. Other reagents were purchased from Sigma Aldrich unless otherwise stated.

As previously [17, 18], we used a high-throughput flow cytometer, the Intellicyt® iQue Screener Plus (Sartorius, Göttingen, Germany), with the following protocol: buffer equilibration (QSol, Sartorius) and plate shaking at 2000 rpm for 50 sec, sampling for 2 sec with 1 sec upload time, 5 sec wash in Qsol buffer every 3 wells, and further probe wash for 10 sec every 12 wells. The instrument has three LED lasers (405nm, 488nm, 640nm) and collects data for two light scattering and 13 fluorescence channels for all samples. Data were analysed and displayed using a combination of the instrument’s Forecyt™ software, FlowJo, and routines written by the first author in R. In contrast to some instruments [44] (and as is evident from the data), this instrument is highly resistant to the detection of extracellular fluorescence. As with any other transport assay method such as those based on filtration or on flow dialysis [45], we do not directly discriminate molecules that are bound from those that are intracellular; other arguments given below serve to do that.

For the dose-response regression analyses the data were fitted with a non-parametric approach (locally weighted scatterplot smoothing) as implemented in R’s ggplot2. The hierarchical dendrogram and heatmap of the palette of 39 fluorophores were scripted with R’s *dendextend* and *ggplot*’s *heatmap.2* packages. The *hclust* function of *dendextend* produced hierarchical clusters from on a Tanimoto similarity matrix derived as follows: fingerprints of 39 fluorophores were derived from their SMILES using the Patterned algorithm within the RDkit (www.rdkit.org/) nodes in KNIME (http://knime.org/) [46, 47]; a Tanimoto distance (TD) matrix was generated from these fingerprints (KNIME’s Distance Matrix Calculate); and the scores of this matrix were converted to Tanimoto similarity indices (1-TD) using KNIME’s Similarity Search. In one case, the hierarchical dendrogram was circularised with R’s *circlize* package to allow space for the composite with the structures of these 39 fluorophores.

## Results

### Baseline analysis

A total of 143 fluorophores were tested in *E. coli* BW25113 (Supplementary Table 1 and Supplementary Figure 1). Figure 1 shows typical cytograms of stained and unstained cells at four concentrations of two particular dyes, in this case calcein (A) and fluorescein (B). The typical existence of a small population with a very high fluorescence biases the mean and so we normally use the median fluorescence for data analysis. The median fluorescence across the entire set of fluorophores spans some three orders of magnitude. Figure 2A shows the distribution of the median fluorescence for each of the dyes at various concentrations. Autofluorescence (the fluorescence from cells to which no dye has been added) provides a ‘baseline’ above which any fluorescence resulting from dye uptake must be resolved (though note that some dyes can act to quench autofluorescence). The red vertical dotted line (Figure 2A) shows the overall median fluorescence when autofluorescence is included while the blue dotted line is the mean fluorescence of dyed wild-type *E. coli* for the 13 fluorescence channels in our Intellicyt^®^ instrument. Channel VL2 exhibits the highest values of autofluorescence relative to the other channels (Figure 2B). Also, a fractional carry over between wells was observed in one of the red channels. The latter is observed as a separate cluster at the very highest values in channel RL1. To avoid this, future experiments were designed, and the data filtered, accordingly. Figures 2C and 2D show the same data after correction for autofluorescence. (Note that because dyes can potentially quench autofluorescence, the fluorescence changes induced by dyes include such phenomena.)

**Figure 1.**
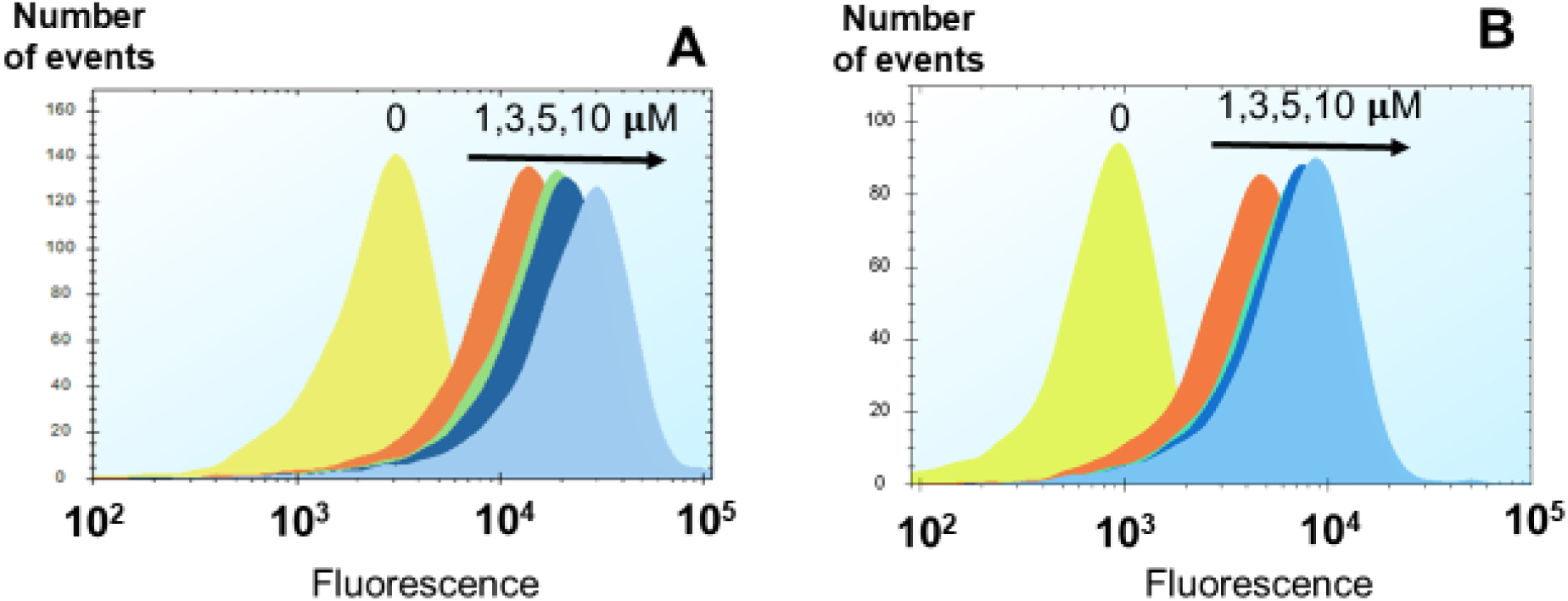
Cytograms of Calcein and Fluorescein. Cells were grown, harvested and re-inoculated at a concentration as described in Methods. Both dyes detected in channel BL3 of Intellicyt® [ex = 488, em = 615/24]. **A**: Calcein fluorescence signals at 1µM, 3µM, 5µM, and 10µM against the non-dye control (0). **B**: Fluorescein signals at 1µM, 3µM, 5µM, and 10µM against the non-dye control (0).

**Figure 2.**
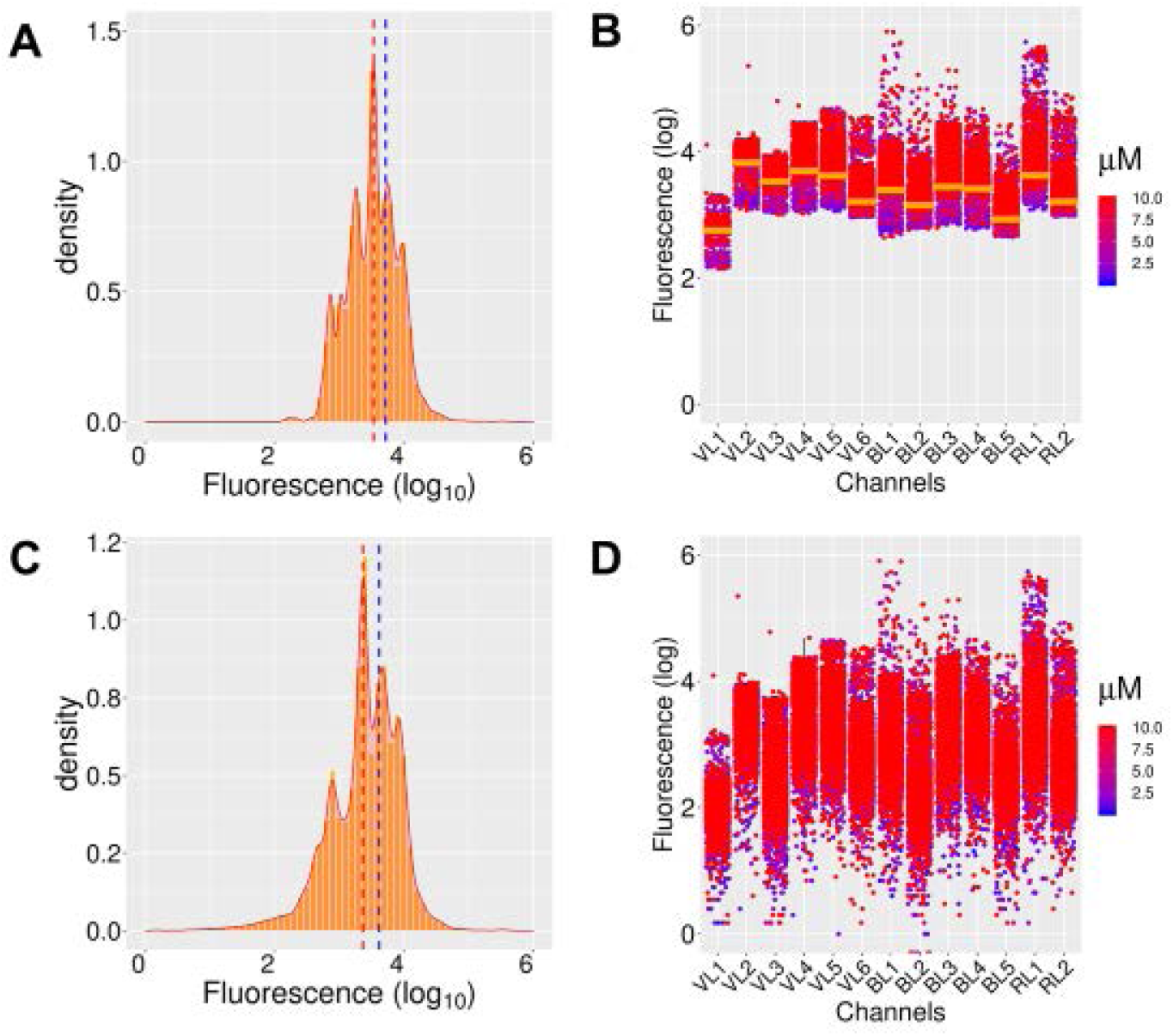
Compilation of the fluorescence detected for 143 fluorescent molecules by the Intellicyt® in *E. coli* BW25113. **A**: Density distribution of the median fluorescence of the full set of fluorophores using their individual optimal excitation-emission channels at their optimal concentrations (as in Supplementary Table 1). **B**: The signals for all fluorophores are compiled against autofluorescence for each channel. Colour-coded distribution of data by concentration of fluorophore (micromolar) are shown. Orange bar: median value of autofluorescence (note that some dyes can quench autofluorescence). **C**: Density distribution after autofluorescence data were subtracted. **D**: The signals for all fluorophores for the 13 channels after subtracting autofluorescence. Colour-coded distribution of data by concentration of fluorophore (micromolar).

A set of 47 fluorophores that showed median values at least twofold (log10 greater than ∼ 0.3) or more than the median of the autofluorescence for a given channel at the concentration used was the core of the set for the next phases of the study (Table 1 and Supplementary Table 1). Inspection of the data showed no particular bias towards either low MW, polarity, nor excitation wavelengths. Two of these dyes are the two that we had used previously [17, 18] (DiSC3(5) and SYBR Green I). The compiled data for four of these molecules (DiSC3(5), pyronin Y, SYBR Green I and Thiazole orange) are shown in Figure 3. They display their strongest signals in channels RL1, BL3, BL1 and BL1, respectively. The set of 47 molecules displayed a reasonable dose response, as judged by accumulated fluorescence (ignoring autofluorescence) in the concentration range 0.1 to 10µM (Supplementary Figure 2). Representative data are given in Figure 4, indicating the tendency to self-quenching for 5-carboxyfluorescein.

**Figure 3:**
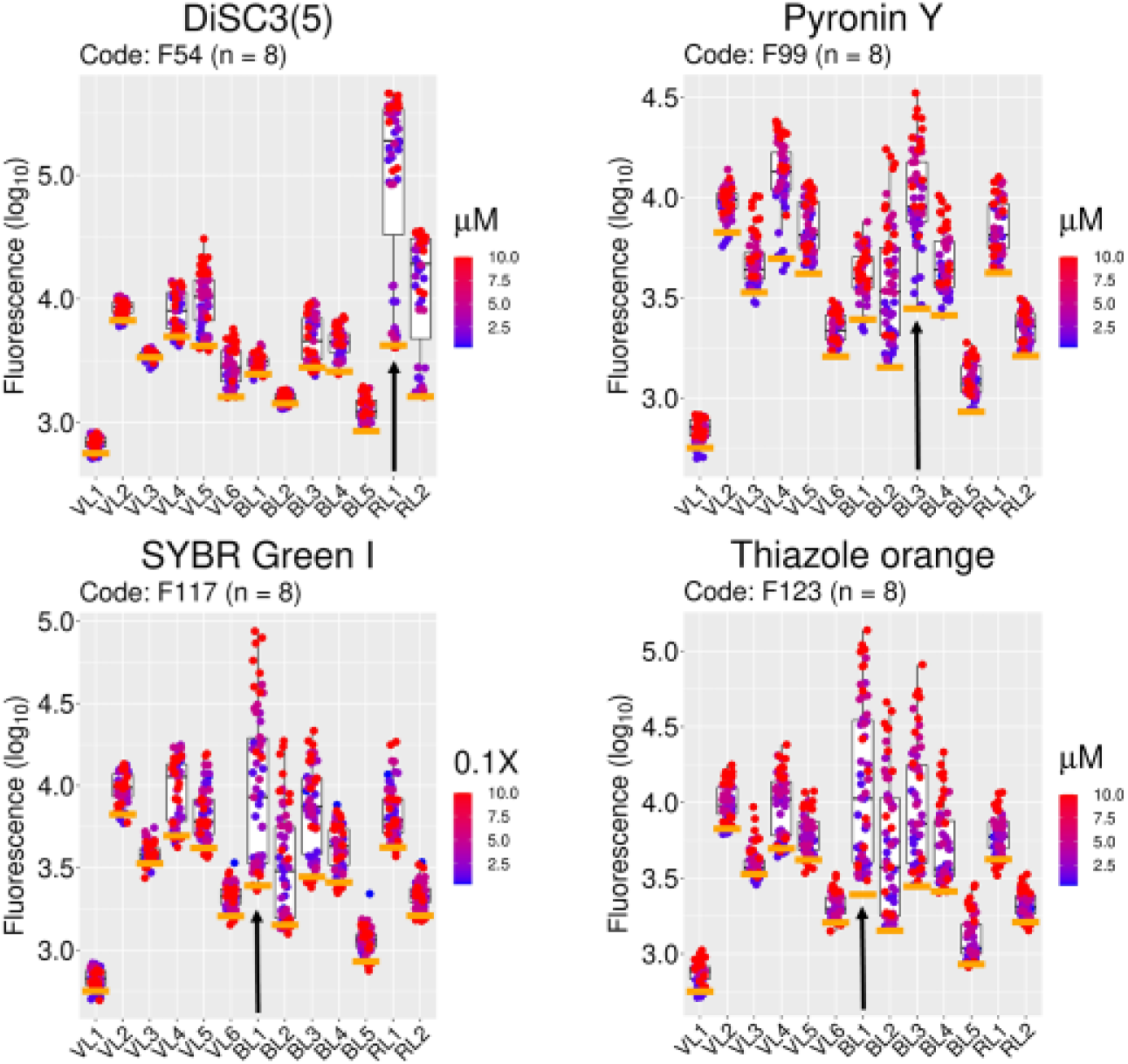
Examples of specific fluorophore uptake in *E. coli* BW25113. Fluorescence signals from four fluorophores selected from the set of 47 molecules (Table 1) with values two-fold or higher above autofluorescence. The signals for all fluorophores (log_10_ values, ordinate) are compiled against autofluorescence (orange bars) for every channel (abscissa). Colour coded distribution of data by concentration of fluorophore (micromolar) are shown. The median values (log_10_) in channel RL1 for DiSC3 and BL3 for Pyronin Y are 50-fold (at 1.3µM) and 4-fold (at 1.3µM) above autofluorescence, respectively. In channel BL1 SYBR Green I (stock diluted 10,000X) and Thiazole orange (10µM) showed log_10_ values above autofluorescence equivalent to 3.2-fold and 4-fold, respectively. The chosen channel for each fluorophore is mapped with an arrow. Each concentration is represented by eight biological replicates taken over a period of five months.

**Figure 4:**
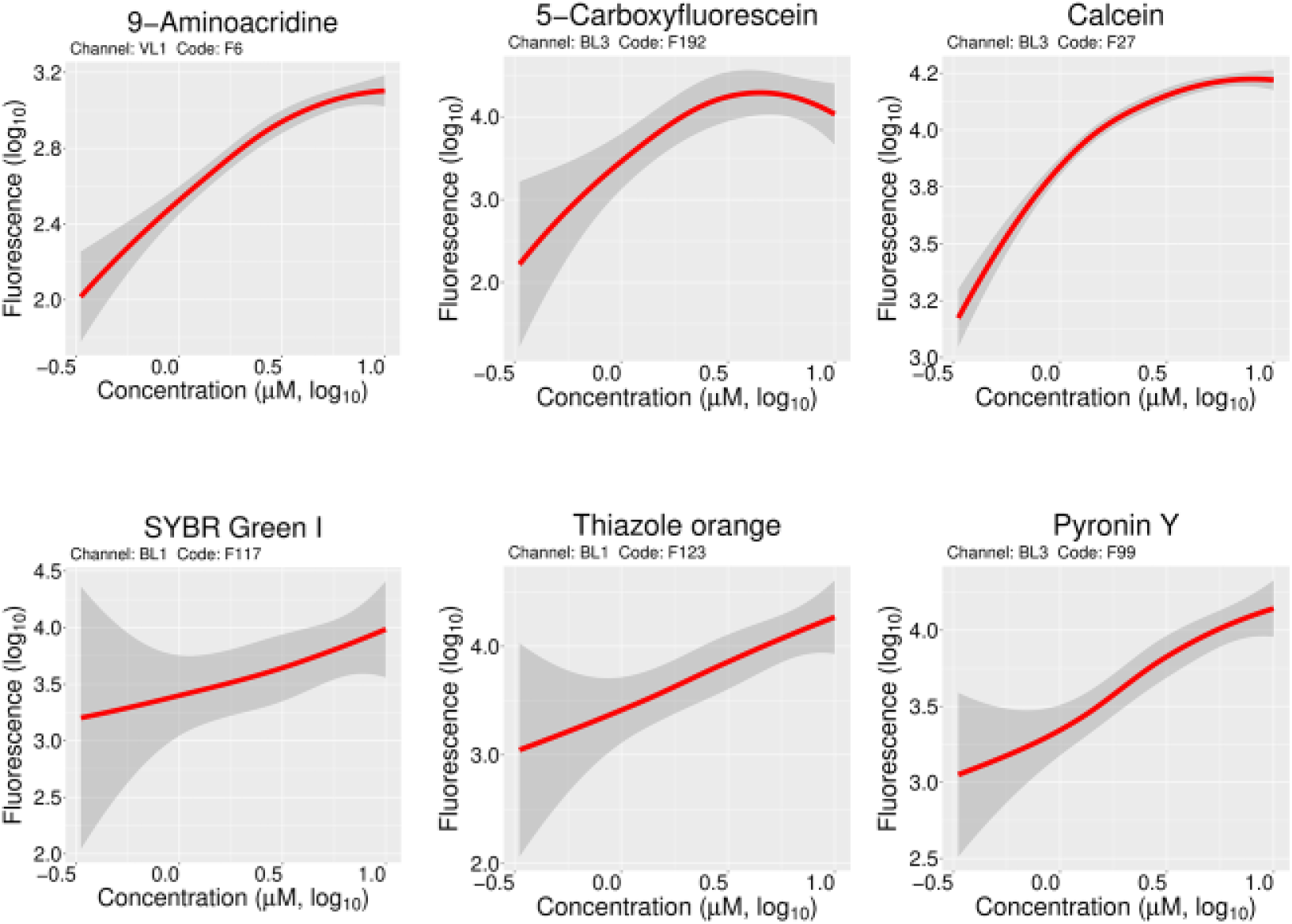
Effect of concentration on fluorophore uptake. Locally weighted scatterplot smoothing was applied to the fluorescence signal versus fluorophore concentration data from six of the 47 chosen fluorophores (see also Supplementary Figure 2). Three fluorophores detected in channels VL1 (9-Aminoacridine) and BL3 (5-Carboxyfluorescein and Calcein) showed linearity only up to 1µM. The dose-response trend was essentially linear for SYBR Green I (channel BL1), Thiazole orange (channel BL1) and Pyronin Y (channel BL3) up to 10µM.

Eight dyes that were quite expensive or of unknown/unavailable structure (i.e., SYTO-13, Cruz Fluor 405, and the six ATTO fluorophores) from the set of 47, were usually not used in follow-up experimentation (except for the gene knockout strain screening shown later). When evaluating the structural similarity of the resultant 39 molecules (Figure 5), a number were phenothiazine derivatives: Acridine orange, Pyronin Y, 9-Aminoacridine, Azure A, Azure B, Azure C, Methylene blue, and Thionine. Xanthene dyes [48] include rhodamines, calcein, fluoresceins and eosin.

**Figure 5:**
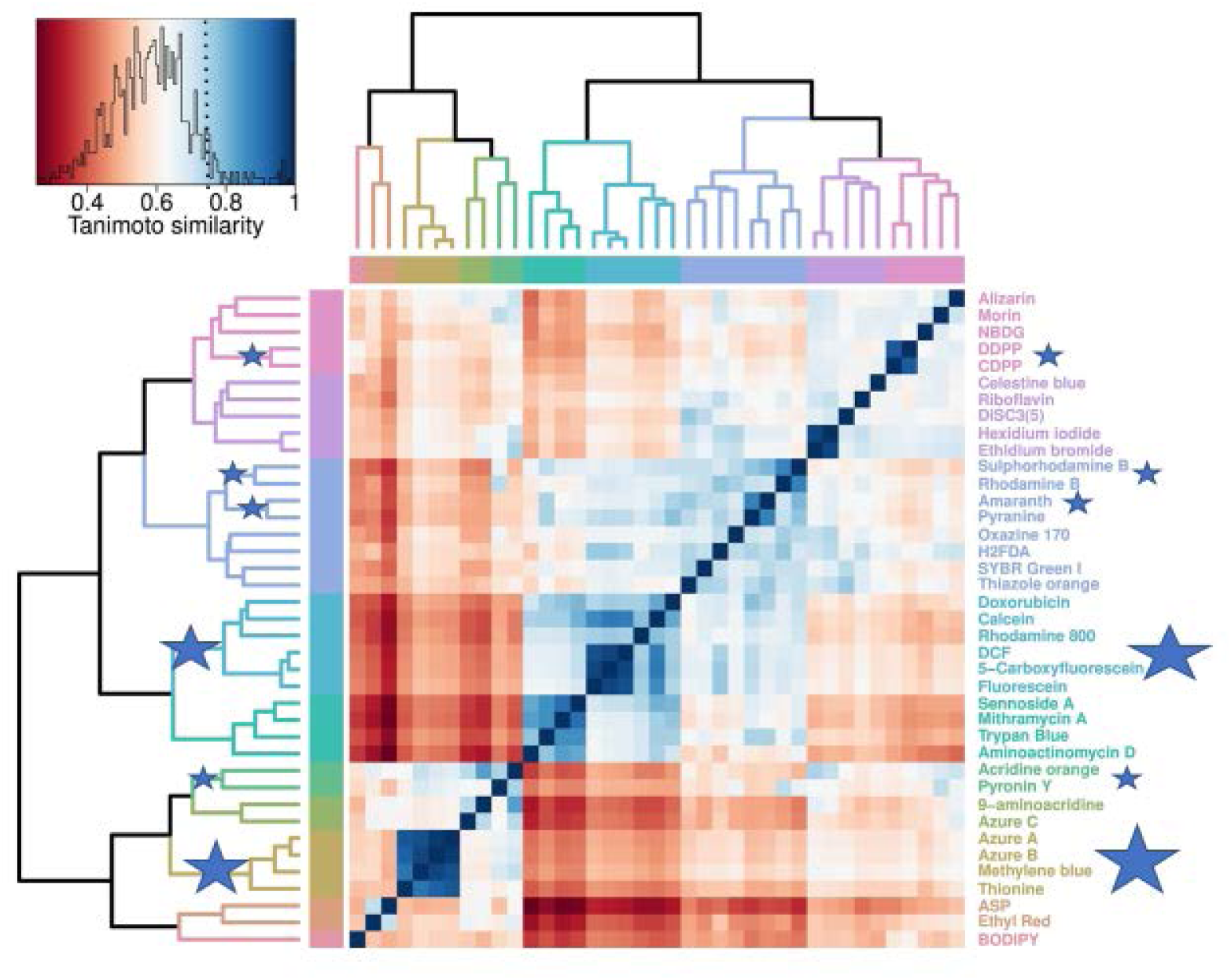
Cheminformatic analysis of fluorophores accumulated by *E. coli* BW25113. Heatmap representing the Tanimoto similarities (derived as described in Methods) of the palette of 39 compounds. Scale from zero (least similarity) to 1 (highest similar). Stars indicate clusters with a Tanimoto similarity exceeding 0.8 (seen as a cutoff for similar bioactivities).

Fluorescein derivatives were the second largest group: Fluorescein, 5-Carboxyfluorescein and 2’,7’-Dichlorofluorescein (DCF), Rhodamine 800, Calcein and Doxorubicin (an antineoplastic antibacterial) (Figures 5). The predicted molecular charges (seen as a decisive property for membrane transport) for these fluorophores showed that NBDG, Acridine orange and Sulphorhodamine B are expected to be zwitterionic at neutral pH (Figure 6). Six other fluorophores are expected to have a net negative charge and 18 to have a positive charge (cationic dyes) (Figure 6). An anticipated example of the latter included the nucleic acid-binding fluorophores (i.e. SYBR Green I, Thiazole orange, Ethidium bromide and Pyronin Y), some of which increase their fluorescence considerably upon such binding. It is of interest that a number are natural products, some with significant molecular masses (e.g. sennoside A, MW 862.7, 7-aminoactinomycin D 1270.4, supplementary table 1), consistent with the view [47] that many transporters evolved and were selected to transport natural products. Pyranine and amaranth are trisulphonated dyes (and they cluster), and how they might get into cells is of some interest.

**Figure 6:**
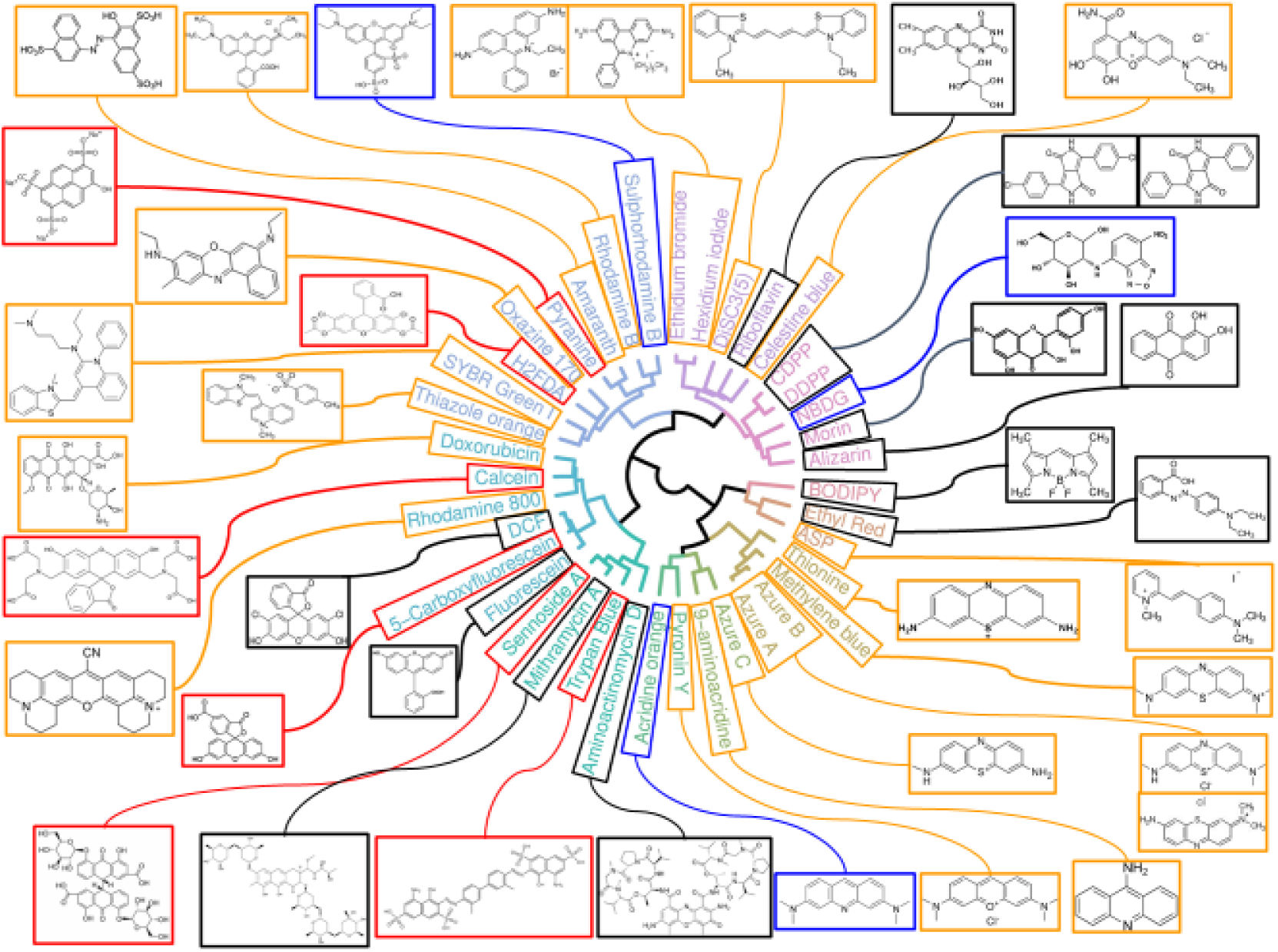
Chemical diversity among 39 fluorophores taken up by *E. coli* BW25113. Circularised topology of the dendrogram in Figure 5 for the visualisation of the structures of this set of fluorophores. The colour of the box indicates the molecular type and/or the expected charge at pH 7.4: black = uncharged, blue = zwitterionic, red = negative, and orange = positive.

Overall, the dyes cover a heterogeneous swathe of chemical space (an overall median Tanimoto similarity below 0.6), and very few form clusters with a Tanimoto similarity greater than 0.75, the lower end of the cutoff region in typical cheminformatic analyses for similar bioactivities [47, 49-52]. The structura heterogeneity is reasonably equated with functional heterogeneity in uptake via different transporters, and this is assessed next.

### Membrane transport mediates intracellular fluorophore accumulation

Most of the fluorophores of interest were taken up rapidly by *E. coli* BW25113, and the intracellular fluorescence as determined flow cytometrically had reached levels within two minutes that did not vary substantially at 15 minutes (Figure 7). A few fluorophores such as DiSC3(5), Oxazine 170, SYBR Green I, and Thiazole orange were accumulated more slowly, but had reached an approximate steady state by the end of 15 minutes.

**Figure 7:**
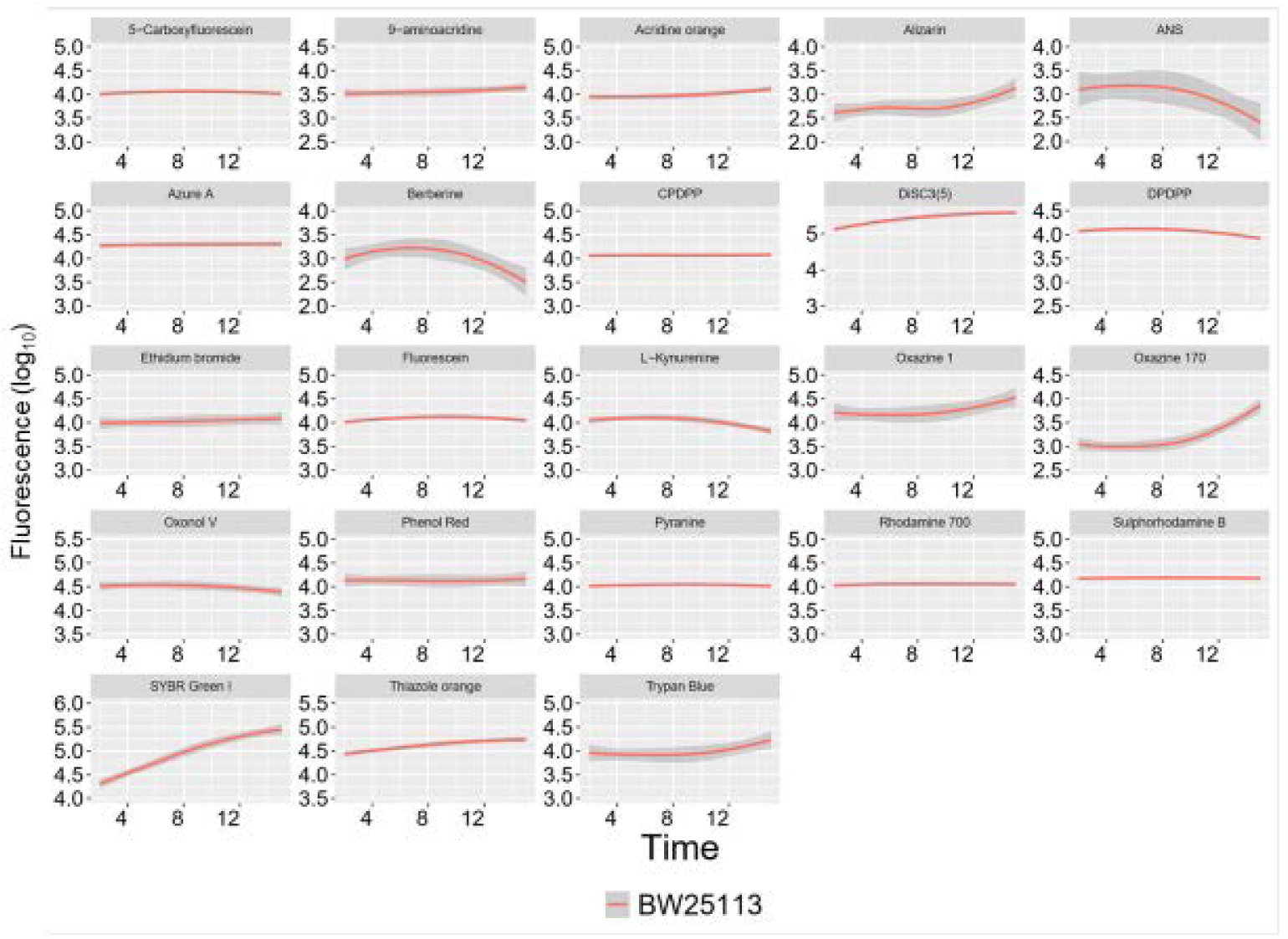
Representative time courses for fluorophore uptake. Locally weighted scatterplot fitting was applied to the fluorescence signal versus time data to follow the levels of fluorophore uptake in the time frame relevant to the experimental settings. X-axis: Time in minutes (up to 15). Y-axis: the log_10_ of fluorescence signals. For the fluorescence uptake experiments with different incubation times. The subset shown consisted of 23 dyes out of the 39 dyes used earlier.

Although even wild-type strains of *E. coli* are not particularly sensitive to the uncoupler CCCP [53], reducing the extent of membrane energisation using it at 10 μM reduced significantly (by more than ten-fold) the uptake of DiSC3(5) into *E. coli* BW25113, while a similar, although smaller effect was observed for thiazole orange, SYBR Green I and pyronin Y (data not shown). Another important parameter that can affect membrane transport is external pH. The accumulation of all fluorophores tested could vary somewhat with pH in the range tested (pH 6 – 8.5), with the highest effect (up to two orders of magnitude) seen in the DPP-based dyes CPDPP and DPDPP (Figure 8). For these two molecules the reduction in their signals is likely due in part to the quenching of their fluorescence because of the deprotonation of the lactam nitrogen in the DPP core [54]. The pH-related behaviour of other fluorophores is presented in Supplementary Figure 3.

**Figure 8:**
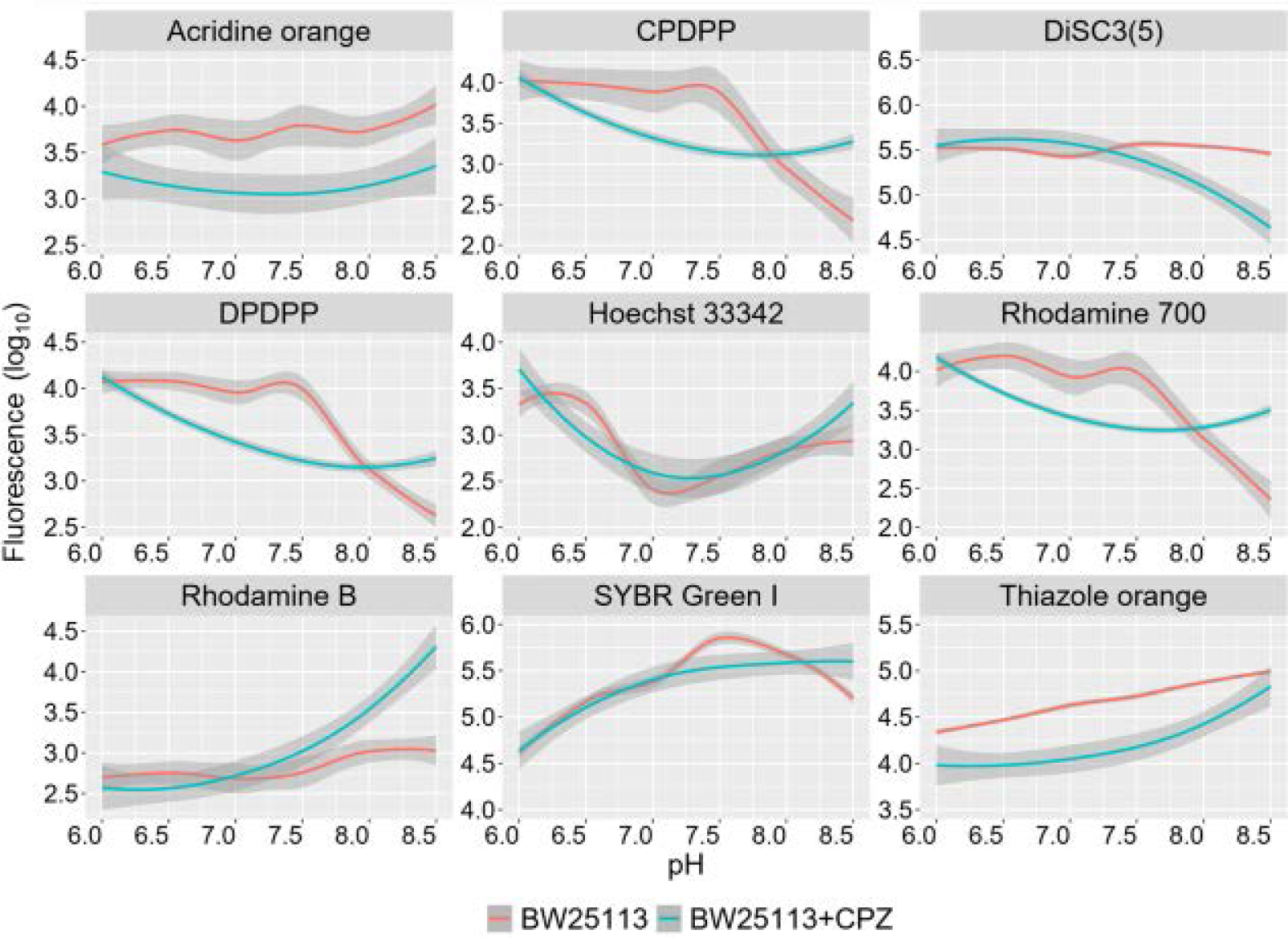
Effect of pH and CPZ on the uptake of various fluorophores. *E. coli* BW25113 (10^6^ cells.mL^-1^) were incubated with 3µM fluorophores for 15 minutes at 37°C (red lines). Similar samples were incubated with both 3µM fluorophores and 10µM CPZ under the same experimental conditions (blue lines). The incubations were carried out at different pH values: 6.0, 6.5, 7.0, 7.5, 8.0, and 8.5. X-axis: pH values. Y-axis: log10 of fluorescence signals after locally weighted scatterplot smoothing. The effect of pH and CPZ on the rest of the palette of fluorophores transported by *E. coli* is given in Supplementary Figure 3.

Chlorpromazine is a known efflux inhibitor in *E. coli* [18, 55-57]. In strain BW25113, CPZ increased the accumulation of Rhodamine B by an order of magnitude and that of SYBR Green by four- or five-fold (Figure 9). The fluorescence signal from Oxazine 1, Oxazine 170 and ASP (4-(4-(Dimethylamino)styryl-N-methylpyridinium (ASP+)) was increased three-fold. Here, however, the opposite effect was shown for DiSC3(5), with CPZ inducing a decreased uptake of some three-fold. We previously published significantly greater effects of CPZ on SYBR Green I uptake in *E. coli* MG1655 [18] (almost a 20-fold increase). We repeated these experiments for strain MG1655, with the same results as previously: the explanation for the difference with strain BW25113 is that BW25113 can accumulate far more di-SC3(5) in the absence of CPZ than does strain MG1655 under the same conditions. The uptake in the presence of CPZ is fairly similar for the two strains, implying a much lowered basal expression of efflux pumps such as acrAB/tolC in strain BW32113 (Figure 10A). The breadth of the distribution of uptake is consistent with this. Such data illustrate the potential for very substantial variation in the uptake of individual dyes between strains of the same organism. Furthermore, we observed significant differences in the light scattering of these two strains of *E. coli* (Figure 11B), especially in the forward scattering (which reflects differences in cell size distribution [7, 58]).

**Figure 9:**
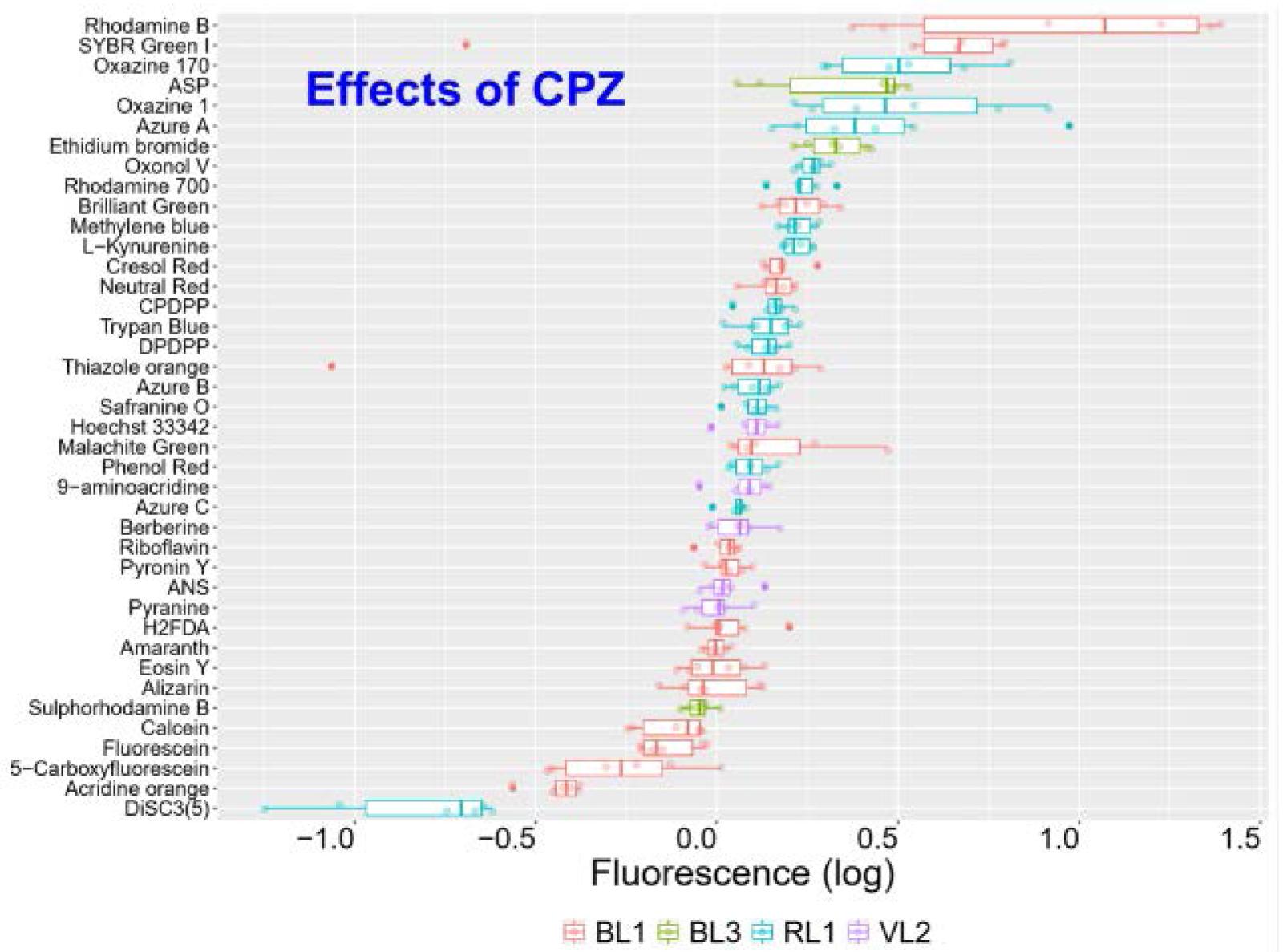
Effect of CPZ in the uptake of fluorophores in LB at pH 8.5. Fluorescence signals are presented as the log_10_ of ratios of the fluorescence from treated cells (*E. coli* BW25113 and 10µM CPZ) over the fluorescence signals from untreated cells. This time the fluorophore uptake incubation (37°C, 15 min) was carried out at pH 8.5, which showed the strongest deviations from a ratio of 1 (log_10_ = 0). Boxplots are ordered by median values. Colours encode the IntelliCyt fluorescence channels used.

**Figure 10:**
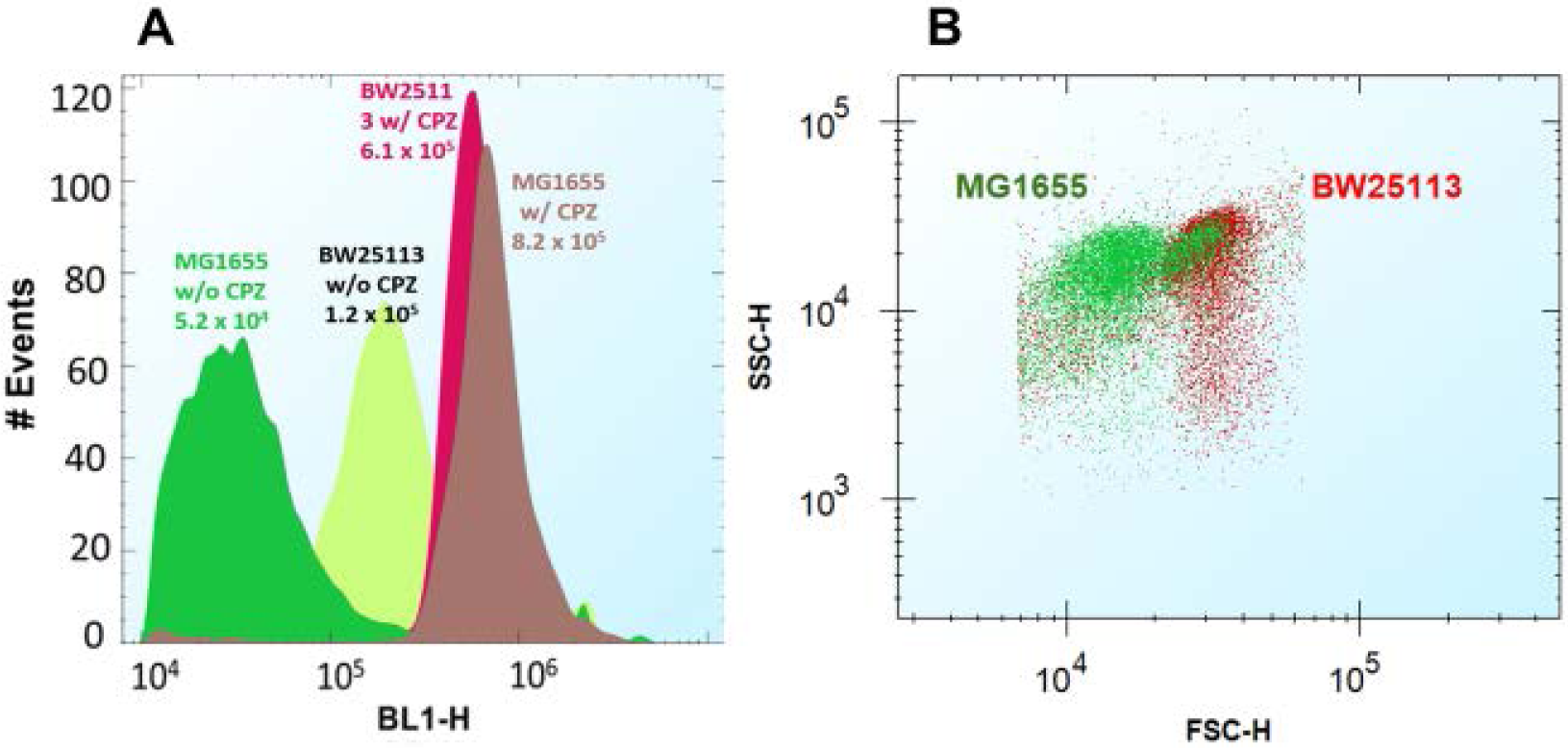
Difference in fluorophore accumulation and light scattering between *E. coli* strains MG1655 and BW25113. **A**: Cytograms showing the effects of CPZ on the uptake of SYBR Green into the two strains. The experiments were performed using the same settings published in [18], as also mentioned in Materials and Methods. **B**: Dot plots of light scattering for these two strains. FSC-H: forward scattering and SSC-H: side scattering. Both strains were processed identically and in parallel for these fluorophore uptake assays.

**Figure 11:**
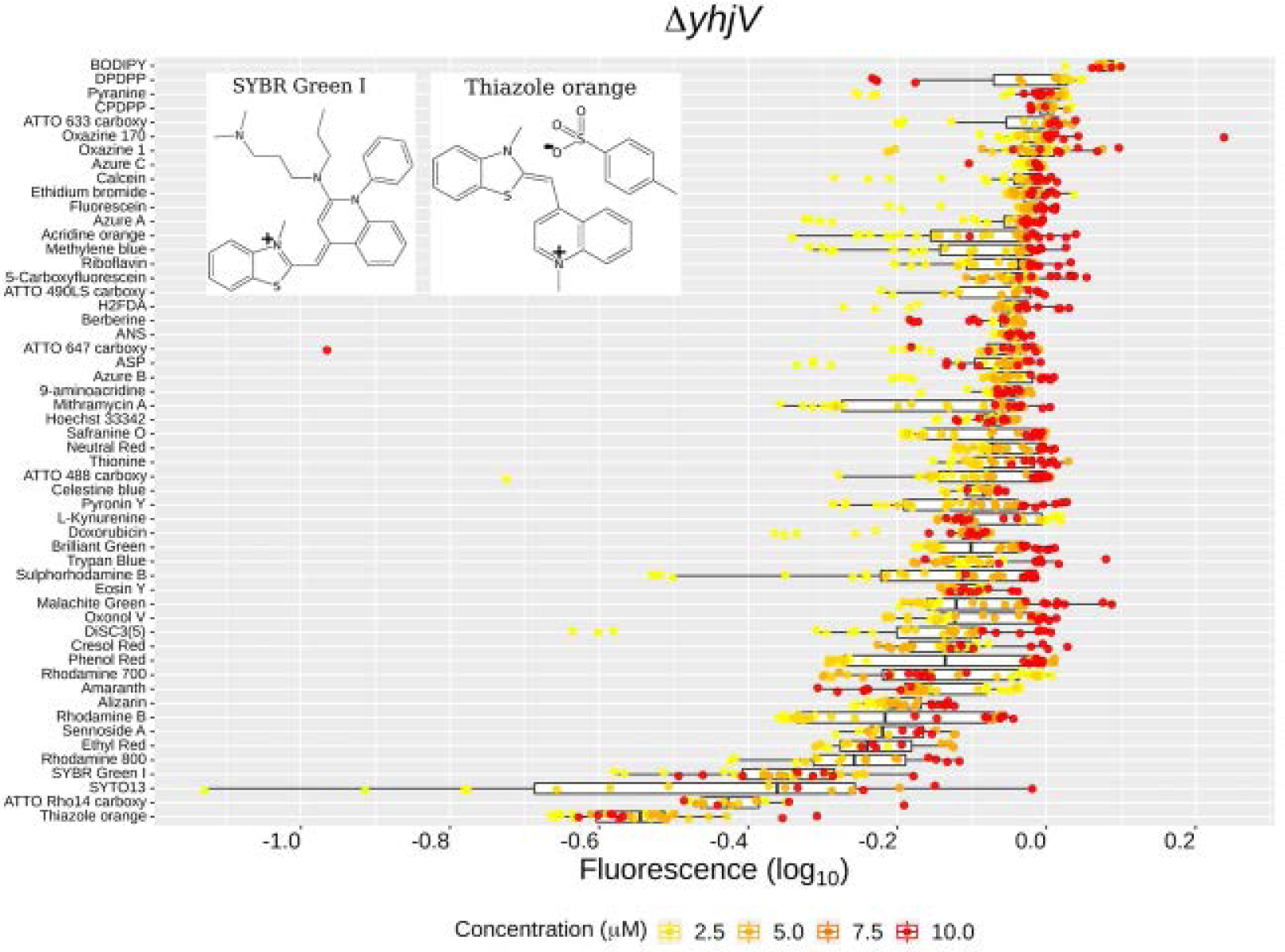
Effect of *yhjV* deletion on fluorophore uptake in *E. coli* BW25113. Fluorescence signals are presented as the log_10_ of ratios of the fluorescence from the *yhjV* knock out cells (Δ*yhjV*) over the fluorescence signals from the reference cells (BW25113). X-axis: log_10_ of the ratios (log_10_ = 0). No difference between the two strain would have a ratio of 1 (log_10_ = 0). Boxplots are ordered by median (highest on top of the plot). The legend lists the concentrations used for each fluorophore. The legend lists the concentrations used for each fluorophore. The clearly related structures of the two most differentiating dyes from the palette of 39 are shown.

### Profiling membrane transporters using fluorophores

As indicated, one of our interests is in the deorphanisation of y-gene transporters. An expanded set of fluorophores was used to interrogate the functional traits of strains lacking one of three genes encoding membranes transporters, viz *yhjV, yihN*, and *tolC*. The success of the gene knockouts was confirmed by PCR (not shown). Two of these genes, *yhjV* and *yihN*, encode transporters with unknown substrates. The third one, *tolC* encodes an ancillary protein component that is used by a number of different membrane transporters, mainly effluxers. Figures 11 and 12 indicate that, after normalising for autofluorescence, the y-gene knock outs Δ*yhjV* and Δ*yihN* showed a clear “influxer” pattern of fluorescence whereby the median signals of the tested fluorophores were below that of the reference strain (i.e., below a ratio of 1, log10 = 0). For the majority of dyes the uptake responded to concentrations of up to 10 µM. The fluorescence signal ratios of KO strains versus the reference strains are given in Table 2 for all three strains, together with their standard deviations and the 95% confidence interval.

**Figure 12:**
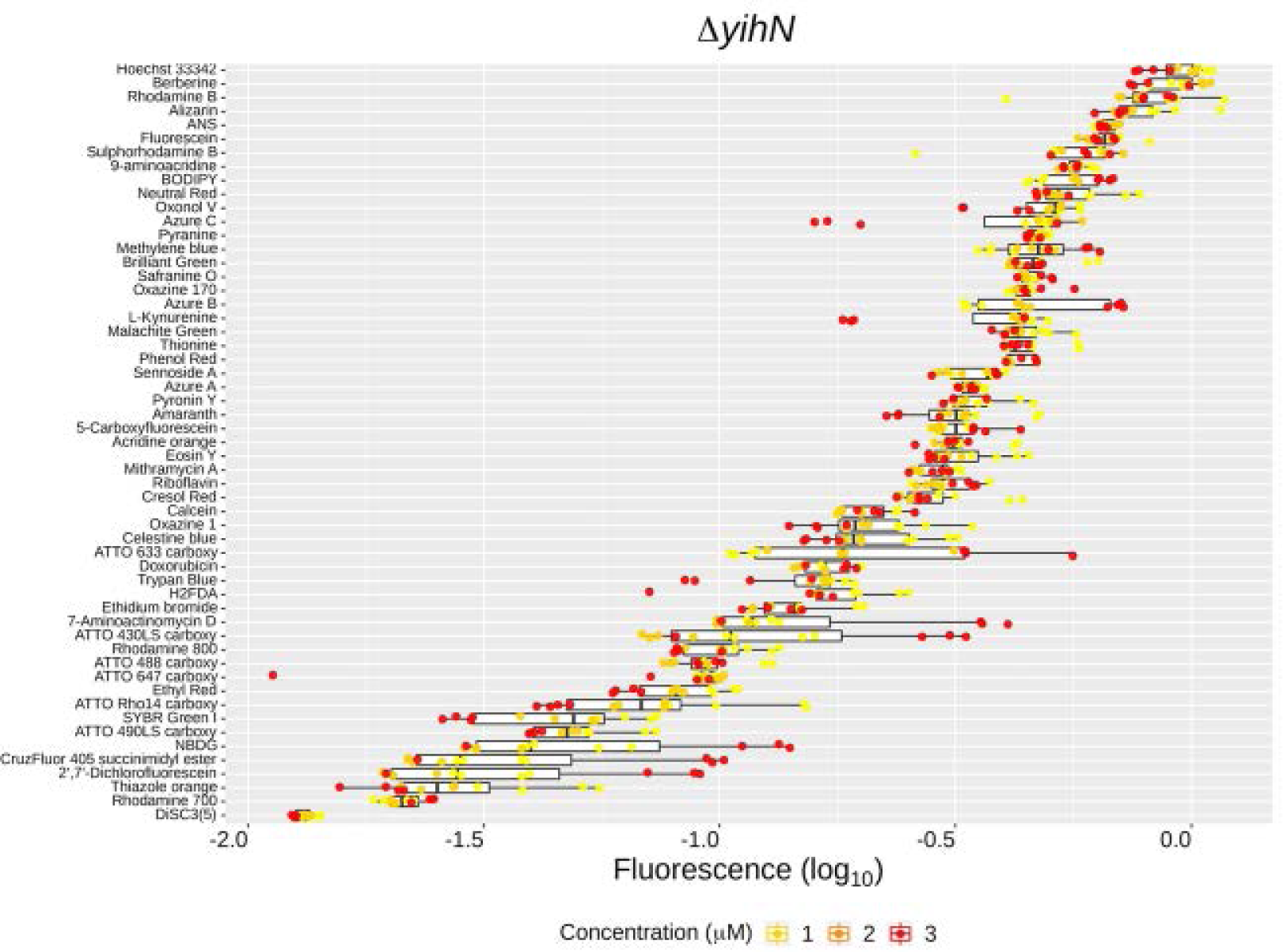
Fluorophore uptake in the *E. coli* gene knock out strain *yihN*. Fluorescence signals are presented as the log_10_ of ratios of the fluorescence from the *yihN* knock out cells (Δ*yihN*) over the fluorescence signals from the reference cells (BW25113). X-axis: log_10_ of the ratios (log_10_ = 0). No difference between the two strain would have a ratio of 1 (log_10_ = 0). Boxplots are ordered by median (highest on top of the plot).

All fluorophores had a reduced uptake in Δ*yihN* with most of them accumulating less than three-fold the amount in comparison to *E. coli* BW25113 (Figure 12). Both Rhodamine 700 and DiSC3(5) showed the lowest uptake for any of the concentrations tested (up to 3 µM). Thiazole orange and SYBR Green I were also poorly accumulated by Δ*yihN*, with more than a ten-fold difference relative to the reference strain (Figure 12, Table 2).

*E. coli* contains a great many ‘efflux’ pumps, mutations in which can cause substantial resistance to multiple drugs and antibiotics (e.g. [59-65]). Many are driven by ATP hydrolysis [66] though others obtain free energy via electron-transport-linked membrane energisation. A variety of antiporters can also, under some circumstances, appear to act as ‘effluxers’ [30, 67, 68]. TolC is an outer membrane component that interacts with many inner-membrane-located transporters (exemplified by acrAB) [61, 69-74]. The involvement of tolC with efflux systems was evident in the pattern of fluorophore uptake observed for 56 dyes in this knockout strain (Figure 13, Table 2). Most fluorophores were over-accumulated (ratios over 1, or log10 > 0). Five fluorophores over accumulated mainly at the higher concentration of 10 µM: fluorescent glucose (NBDG), 2’,7’-Dichlorofluorescein, 7-Aminoactinomycin D, ATTO 430LS, and CruzFluor 450. On the other hand, nine fluorophores had a decreased accumulation of two-fold or lower in Δ*tolC*, with a ten-fold or lower differences in the cases of Rhodamine 800, DiSC3(5) and Rhodamine 700 (Figure 13, Table 2). That the lack of *tolC* seems to impair the transport of the latter dyes points at the interaction or involvement of this protein with outer membrane <u>influx</u> systems (e.g., porins), either directly or via pleiotropic effects [75, 76]. The data for the three knockout strains and the 39 dyes are shown in Fig 14 (with the six major dyes for the yihN knockout being labelled, and with all the data being given in Table 2).

**Figure 13:**
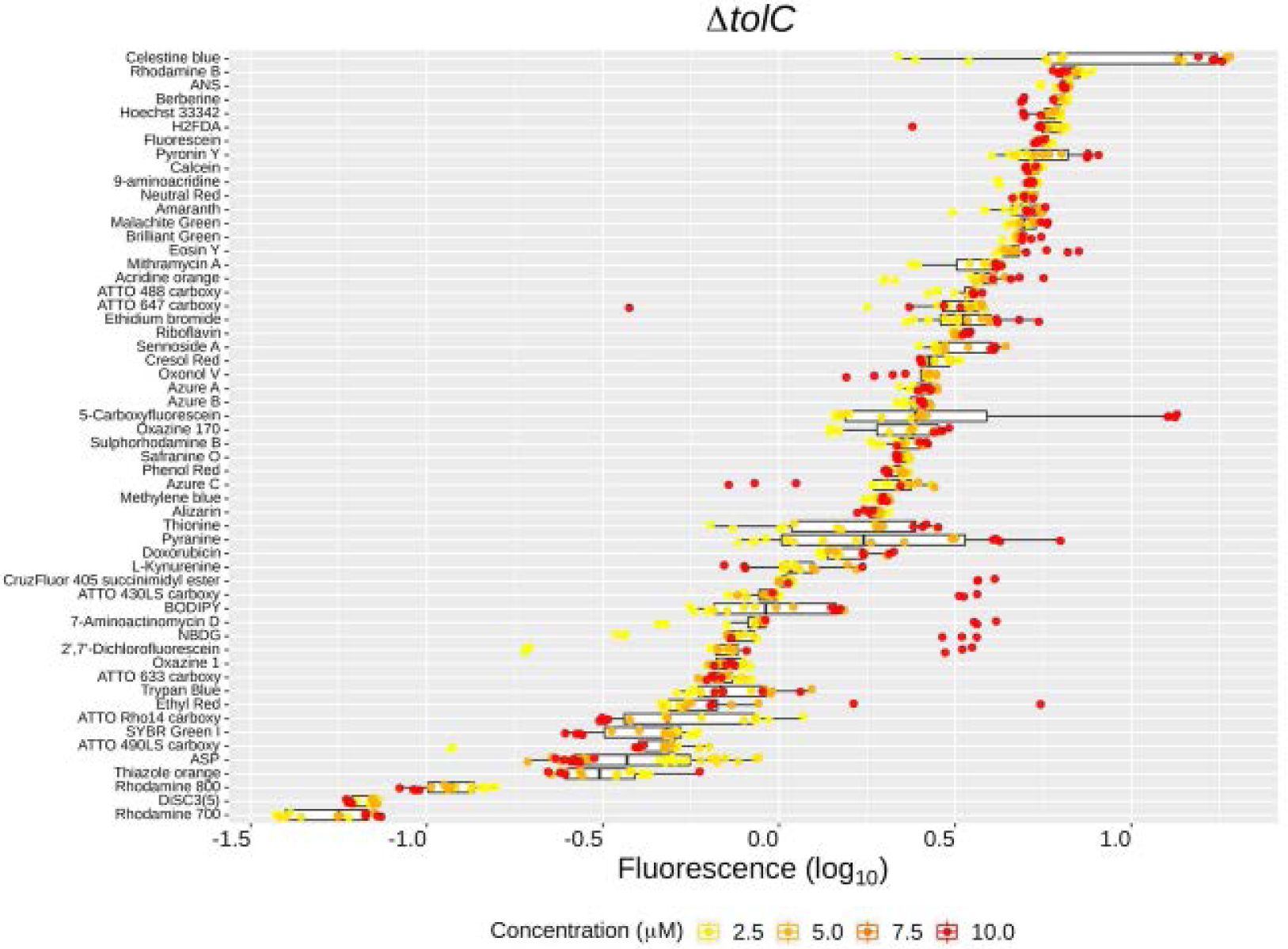
Fluorophore uptake in the *E. coli* gene knock out strain *tolC*. Fluorescence signals are presented as the log_10_ of ratios of the fluorescence from the *tolC* knock out cells (Δ*tolC*) over the fluorescence signals from the reference cells (BW25113). X-axis: log_10_ of the ratios (log_10_ = 0). No difference between the two strain would have a ratio of 1 (log_10_ = 0). Boxplots are ordered by median (highest on top of the plot). The legend lists the concentrations used for each fluorophore.

**Figure 14:**
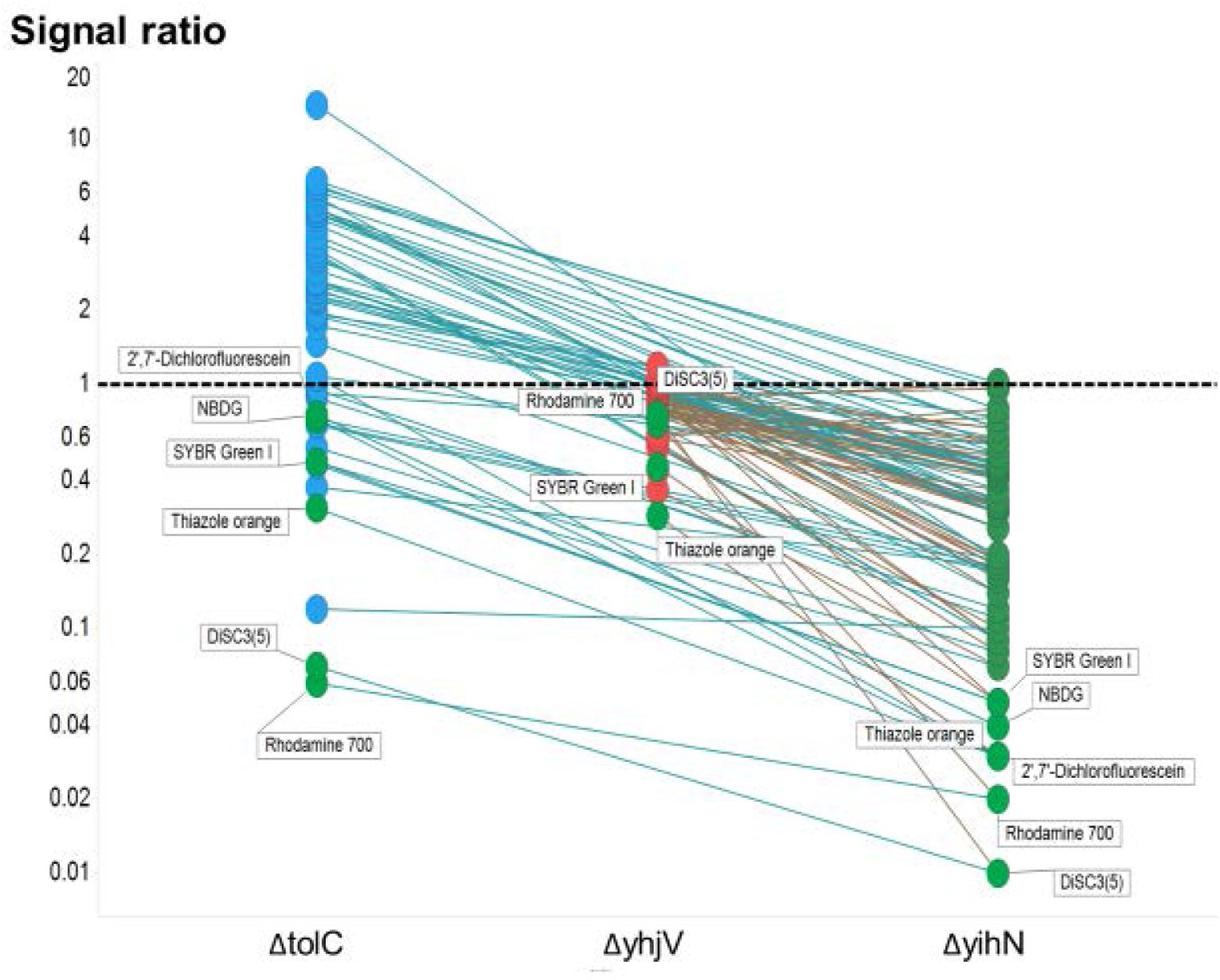
Differential uptake of 39 dyes between three different knockout strain of *E. coli*. Experiments were performed as described in the legends to figures 11-13 and in Table 2. Several dyes showing the largest effect in ΔyihN are marked.

Finally, we also considered the opposite kind of experiment, in which specific transporter genes are carefully overexpressed, using the equivalent ASKA overexpression clones [77, 78]. Figure 15 shows the cytograms for the uptake of thiazole orange into the deletant, wild-type and overexpression strains for yhjV when the concentration of the dye was 1 μM, while Figure 16 shows more extensive data for three dyes at four concentrations. Each of these dyes is clearly capable of providing a surrogate transport assay for the yhjV transporter.

**Figure 15:**
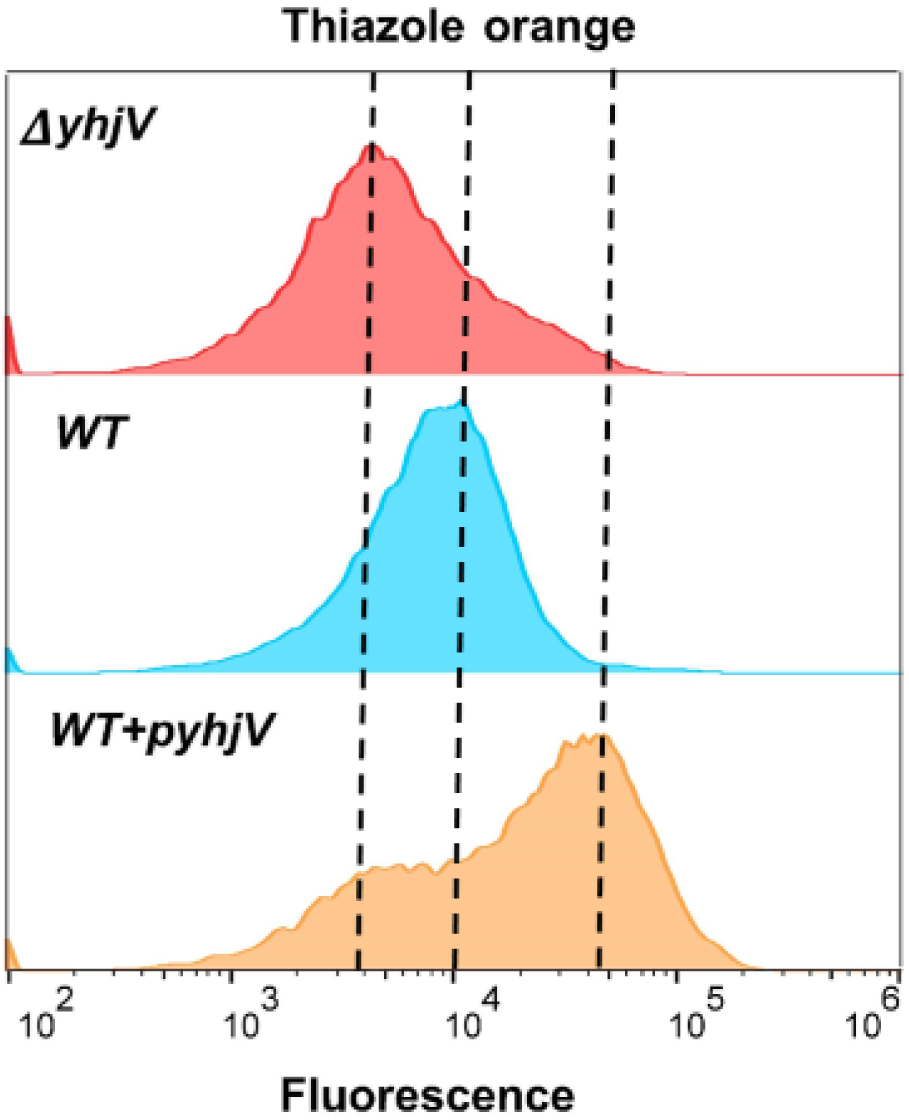
Cytograms of the uptake of thiazole orange by *E. coli* BW25113 with differing expression levels of *yhjV*. The cell population distribution for the accumulation of thiazole orange (1µM, 37°C, 15 min) is presented for the gene knockout of *yhjV* (Δ*yhjV*) of the Keio collection. The fluorescence signal of reference strain *E. coli* BW25113 (*WT*) is approximately 2.5-fold higher than the signal from the gene knockout strain (Δ*yhjV*). The strain of *E. coli* expressing the *yhjV* gene from the ASKA collection (*WT+pyhjV*) shows a further increase in the accumulation of thiazole orange equivalent to approximately 4.5-fold over the signal from the WT ((data taken from the assay where the highest differences were observed for 1µM).

**Figure 16:**
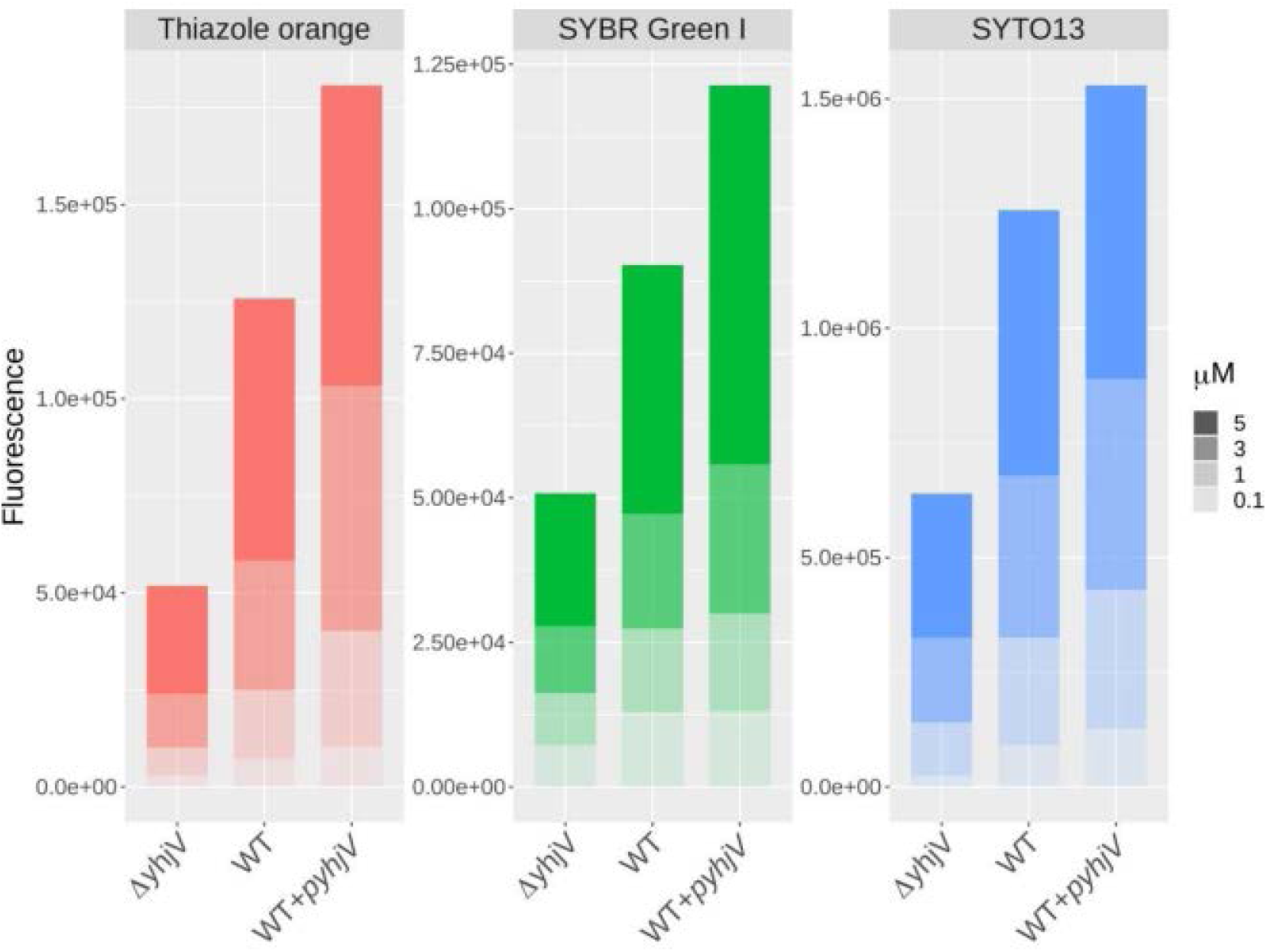
*E. coli* BW25113 overexpressing *yhjV* displays increased uptake of thiazole orange, SYBR Green I and SYTO13. The fluorescence signals of cells accumulating these fluorophores (37°C, 15 min) are presented for the gene knockout for *yhjV* (Δ*yhjV*), the reference strain *E. coli* BW25113 (*WT*) and the strain overexpressing the ASKA construct of *yhjV* (WT+*pyhjV*). Cells were exposed to four different concentrations of fluorophores up to 5µM, as indicated. Fluorescence data are the averages of the median values from four biological replicates. The accumulation of these fluorophores in all three strains increased monotonically with ascending dye concentrations. For each Fluorophore WT cells gave higher fluorescence signals than did strains deleted in yhjV, and likewise cells expressing episomal yhjV (WT+pyhjV) over the reference strain (WT). The latter relation for Thiazole orange at 1µM is somewhat smaller (approximately 2.62-fold) than that shown in figure 15 (4.5-fold) since here we have the average of four experimental replicates.

## Discussion

### Role of fluorescent stains in understanding microbial physiology

Fluorescent molecules continue to be instrumental in the versatile, sensitive and quantitative study of microbial structure and physiology. However, the use of fluorophores in the characterisation of membrane transporters continues to be sparse (it is done mainly in mammalian cell lines) and is largely directed to the study of drug resistance mediated by efflux pumps (e.g. [34, 79, 80]). Indeed, many drugs show structural similarities with fluorophores [81]. Early evidence showed that fluorescent dyes such as rhodamine derivatives used in the study of drug efflux require mediated membrane transport influx in the first place [82], and the same is true for nucleic acid stains such as ethidium bromide [83, 84].

Previously, we studied just diSC3(5) [85, 86], which is subject to quenching at high concentrations [87], and SYBR Green I [9, 88-94], where quenching is intramolecular and is stopped (and fluorescence massively enhanced) by binding mainly to dsDNA [95].

Here we have surveyed a large set of fluorophores to assess their suitability as probes for the study of mediated membrane transport in bacteria, initially using the reference strain BW25113 of the Keio *E. coli* gene knockout collection [19].

The desirable properties of such dyes were given in the introduction, and are not repeated here. We do note two caveats, however: (i) that the excitations of those dyes surveyed somewhat reflected the wavelengths of the lasers that we used, and (ii) the eventual choice was largely based on the properties of the wild-type strain. Thus other dyes might have been included if other excitation wavelengths were available, and/or if other strains showing lower expression of (or lacking) particular efflux pumps had been used.

Out of 143 fluorophores surveyed, the median values of 47 provided signals significantly above autofluorescence to consider pursuing. If those with mean values just two-fold above autofluorescence are included this set can be larger (60 fluorophores). However, the latter group of fluorophores are mainly accumulated by the population of higher uptake cells in the right-skewed distributions of fluorescence (Figure 2C). The accumulation of a core of 39 fluorophores was of particular interest, and these were further characterised after selecting against those of higher cost or with unknown structures. These dyes stained the wild-type BW25113 *E. coli* on short time scales (Figures 4, 7) in a manner that depended suitably on substrate concentration (Figures 3,4). Their signals were mostly sensitive to membrane efflux transport inhibitors such as chlorpromazine, and in some cases to the pH of incubation (Figure 8).

### Cheminformatics of chosen fluorophores

Despite the reasonable number of available fluorophores, many are not particularly cheap, and our survey was largely confined to the more inexpensive ones available commercially. Cheminformatics [96-98] (sometimes called chemoinformatics [96, 99-103]) describes the discipline that helps researchers assess questions such as the degrees of similarity between individual molecules [104-106] or the molecular diversity within a library [107, 108]. We applied standard cheminformatics methods [46, 47, 109, 110] to the analysis of the relative diversity of our palette of 39 dyes. Given that a pairwise Tanimoto similarity below 0.8 (or a Tanimoto difference exceeding 0.2) is usually taken to mean a significant difference in bioactivity [47, 49-52], it is encouraging that while there were some small clusters (Figures 5, 6), the median Tanimoto similarity was just 0.6, implying strong orthogonality in the behaviour of our palette, as was borne out experimentally.

This said, many of the diverse dyes were still either phenothiazines or xanthene family dyes, and one conclusion is that the need for a much greater variety of fluorescent scaffolds remains, in order to broaden the present collection yet further.

### Strain differences

It is well known that strain differences between even non-pathogenic *E. coli* can have massive effects even on simple traits such as recombinant protein production [111]. In the present work, in some cases, we observed quite striking differences between the uptake of particular dyes into the two wild-type strains BW25113 and MG1655. In particular, the observed differences in uptake of SYBR Green I in *E. coli* MG1655 and BW25113 might be due to their known genomic differences: *E. coli* BW25113 is Δ(*araD-araB)567* Δ*(rhaD-rhaB)568 ΔlacZ4787* (::rrnB-3) *hsdR514 rph-1*, with the deletion of *araBAD* and *rhaDAB* and the replacement of a section of *lacZ* with four tandem *rrnB* terminators as well as a frameshift mutation in *hsdR* resulting in a premature translation stop codon. *E. coli* BW25113 also contains the *lacI+* allele and not *lacIq*. Other known differences found from genome sequencing are the presence of the *rph-1* allele as well as 20 substitutions and 11 indels [112]. Some of those substitutions directly affect membrane transport systems: *nagE* (N-acetyl glucosamine specific PTS enzyme IIC, IIB, and IIA components), *gatC* (subunit of galactitol PTS permease), *btuB* (vitamin B12/cobalamin outer membrane transporter). Furthermore, any single or combined genomic difference can have a number of pleiotropic cellular effects that could easily account for broad phenotypic differences between these two strains of *E. coli* [113, 114]. This is a typical issue of complex systems biology that can cause difficulties yet also has blessings: the difficulties can occur because of an ostensible non-reproducibility of experiments that are in fact different (but in unknown ways), while the blessing is that it shows that our palette of dyes is a particularly strong discriminator of the physiology of different, and even closely related strains. Having previously seen that even single-gene knockouts could induce massive and completely uncorrelated effects in the uptake of particular dyes [18], we sought to assess the utility of this phenomenon in analysing the general dye uptake properties of three transporters, viz yhjV, yihN, and tolC.

### The profiles observed with gene knockouts discriminate influx from efflux systems

It is reasonable that the genetic knockout of an import transporter (including antiporters [30, 68]) will tend to result in lower uptake of members of the palette than do knockouts of predominantly efflux transporters, that may be expected to have the opposite effect. We assessed this expectation using three transporters, with the expected general results.

As an orphan transporter (marked at Uniprot https://www.uniprot.org/uniprot/P37660 as ‘inner membrane transport protein yhjV’, and as a possible ‘amino acid transmembrane transport protein’), we decided to use yhjV as a first test case for exploiting our dyes. It was chosen because in our previous work [18] the KO strain for *yhjV* accumulated one of the lowest amounts of SYBR Green. There is next to no literature on it, however [115]. Although its main substrates are not known, yhjV is considered to be an uncharacterised member of the Hydroxy/Aromatic Amino Acid Permease (HAAAP) Family within the Amino Acid-Polyamine-Organocation (APC) Superfamily [116, 117]. To assess other dyes, we compared our palette in terms of uptake between the wild-type and Δ*yhjV* strains (Figure 11). With the exception of BODIPY, all dyes were taken up less in the knockout than in the wild type, though only a few by less than threefold. Thiazole orange was an even better (more discriminatory) substrate than was SYBR Green. Despite their rather different names, these two dyes are reasonably similar structurally (Figures 6, 11), with both including a benzothiazole moiety linked to dual-ring systems. This similarity in behaviour is consistent with the principle of molecular similarity [104], by which similar molecular structures are expected to possess similar bioactivities.

*yihN* encodes another transporter (https://www.uniprot.org/uniprot/P32135) of unknown function [118, 119]. Here again, essentially no dyes were taken up more in the knockout than in the wild type (Figure 12), implying again that yihN is mainly an influx transporter. In this case, about a dozen dyes were accumulated less than tenfold in the knockout relative to accumulation in the wild type.

tolC is well known to be involved in the efflux of a great many substances because it is linked to a variety of inner-membrane efflux transporters [69, 72, 120-122]. In this case the uptake of the majority of the palette was, as expected, greater than in the wild type in the tolC deletion strain. Rhodamines 700 and 800 are hydrophobic (6-ringed) cations, closely related structurally, and are the most and third-most dyes in terms of lowered uptake in the ΔtolC strain (Figure 13).

Consequently, the effects of a knockout of a putative transporters on the palette give a clear indication as to whether it is mainly an influx or an efflux transporter (Figure 14) (although we note that under the conditions of individual assays we cannot discriminate symporters and antiporters; that requires the use of multiple conditions including the putatively antiported substrates [30]).

The above experiments included solely gene knockout strains in comparison to the behaviour of the wild type strain. Overexpression strains provide an arguably more powerful, and at least complementary, approach to understanding the biology of individual proteins. To this end (Figure 15), we compared the uptake of thiazole orange in the deletant and overexpression starins of yhjV, using thiazole orange, finding an approximately 9-fold variation in uptake of the dye in the two strains, providing a clear use for it as a surrogate dye in assays for this transporter.

### Concluding remarks

The present results and analysis have, for the first time, provided a reasonably comprehensive set of stains for *E. coli*, and have illustrated how sensitive their uptake can be to genotype and physiology. Clearly the same strategy can be applied to other microbes of interest, where different preferred dyes are certain to be found (note that rhodamine 123 [123] and hexidium iodide [124] readily enter intact cells of Gram-positive but not Gram-negative bacteria). The differential uptake of specific dyes in strains knocked out for or overexpressing particular transporters provides a clear means for high-through assay of these transporters and libraries of potential variants [125-127]; this will be the subject of a future communication.

## Abbreviations

ANS: 8-Anilinonaphthalene-1-sulfonic acid
BODIPY: Difluoro{2-[(3,5-dimethyl-2H-pyrrol-2-ylidene-N)methyl]-3,5-dimethyl-1H-pyrrolato-N}boron
CCCP: Carbonyl cyanide m-chlorophenyl hydrazone
CPDPP: 3,6-Bis(4-chlorophenyl)-2,5-dihydropyrrolo[3,4-c]pyrrole-1,4-dione
CPZ: Chlorpromazine
ASP: 4-(4-(Dimethylamino)styryl-N-methylpyridinium (ASP+)
diS-C3(5): 3-Propyl-2-{(1E,3E,5E)-5-(3-propyl-1,3-benzothiazol-2(3H)-ylidene)-1,3-pentadien-1-yl}-1,3-benzothiazol-3-ium iodide
DMSO: Dimethyl sulfoxide
DPDPP: 3,6-Diphenyl-2,5-dihydropyrrolo[3,4-c]pyrrole-1,4-dione
H2FDA: Dihydrofluorescein diacetate
IPTG: Isopropyl β-d-1-thiogalactopyranoside
NBDG: 2-(N-(7-Nitrobenz-2-oxa-1,3-diazol-4-yl)Amino)-2-Deoxyglucose
SYBR Green I: N,N-dimethyl-N′-[(4E)-4-[(3-methyl-1,3-benzothiazol-3-ium-2-yl)methylidene]-1-phenyl-2,3-dihydroquinolin-2-yl]-N′-propylpropane-1,3-diamine

## Acknowledgements

We thank the BBSRC (grants BB/R000093/1 and BB/P009042/1) and the Novo Nordisk Foundation (grant NNF10CC1016517) for financial support. We thank Drs Kate Baker, Lachlan Munro and Lei Yang for useful discussions. The funding bodies had no role nor involvement in the design of the study and collection, analysis, and interpretation of data nor in writing the manuscript.

## Authors’ contributions

JES-S performed most of the experiments. SJ performed some of them. SO’H performed some of the cheminformatics. DBK contributed the overall design of the study and obtained funding for it. All authors contributed to the writing of the manuscript and approved it.

## Legends to Figures

Supplementary Figure 1: *E. coli* BW25113 fluorescence signals for 143 fluorophores. The signals for the non-dye control (Autofluorescence) and all fluorophores (log10 values, Y-axis) are compiled for each of the 13 channels of the Intellicyt® flowcytometer (X-axis). The median for autofluorescence (orange bars) is drawn for every channel. Colour coded distribution of data by concentration of fluorophore (micromolar) are shown. The subtitles contain our internal code (Fxxx). The number of biological replicates (***n***) for each fluorophore at a given concentration represent data from different experiments carried out over a period of five months.

Supplementary Figure 2: Dose-response fluorophore uptake. Plots with regression analysis for each of the initial 47 fluorophores that had a two-fold or higher activity (in terms of median values of uptake) in *E. coli* BW25113. Locally weighted scatterplot smoothing was applied to fluorescence signal versus fluorophore concentration data. X-axis: the log10 of the concentrations in micromolar. Y-axis: the log10 of fluorescence signals. The subtitles contain the relevant channel for each fluorophore and our internal code (Fxxx).

Supplementary Figure 3: Effect of pH and CPZ on the uptake of 40 fluorophores from which the examples in Figure 8 were taken from. *E. coli* BW25113 (10^6^ cells.mL^-1^) were incubated with 3µM fluorophores for 15 minutes at 37°C (red lines). Similar samples were incubated with both 3µM fluorophores and 10µM CPZ under the same experimental conditions (blue lines). The incubations were carried out at different pH values: 6.0, 6.5, 7.0, 7.5, 8.0, and 8.5. X-axis: pH values. Y-axis: log10 of fluorescence signals after locally weighted scatterplot smoothing.

## Tables

Table 1: Fluorophore set specifically accumulated by *E. coli* BW25113. List of fluorescent (Fluor) molecules that recorded fluorescence signals two-fold or higher (Signal strength) than the median values of autofluorescence. SMILES (Simplified Molecular Input Line Entry System [128]). Channel: Intellicyt® flowcytometer channel that registered the highest median for each fluorophore. The excitation and emission profiles of these channels should have matched the excitation and emission wavelengths for those fluorophores (Supplementary Table 1). The structures of some of these fluorophores are unknown or unavailable (NA). Concentration: concentration at which the highest median values were found for each fluorophore in the range tested. SD: standard deviation of the population, CI95: 95% confidence interval for each fluorophore at the given channel.

Table 2: Fluorophore uptake profile for three different membrane transporters of *E. coli* BW25113. Fluorescence (Signal Ratios) data are presented as the ratios of the fluorescence (median) from the three different *E. coli* knockout strains (Δ*yhjV*, Δ*yihN*, and Δ*tolC*) over the fluorescence (median) from the reference strain (*E. coli* BW25113). No difference between a KO strain and the reference would have had a ratio of 1. SD: standard deviation of the population, CI95: 95% confidence interval. The data for each knockout strain are sorted by incremental signal ratios.

Supplementary Table 1: Set of 143 dyes. Dye: common name for each compound. The median-based subset of 47 dyes that had signal two-fold or higher above autofluorescence are highlighted (YES in *E. coli* BW25113 column). The concentration (µM) for each of those selected fluorophores is also listed (Concentration). IUPAC: nomenclature according the International Union of Pure and Applied Chemistry. SMILES (Simplified Molecular Input Line Entry System). Ex: absorbance wavelength. Em: wavelength fluorescence emission. (Ex and Em are literature values.) NA cells denote unknown data. InChI: IUPAC International Chemical Identifier.

