## Supplementary figures and images for "A palette of fluorophores that are differentially accumulated by wild-type and mutant strains of *Escherichia coli*: surrogate ligands for bacterial membrane transporters"

### Supplementary Fig 1

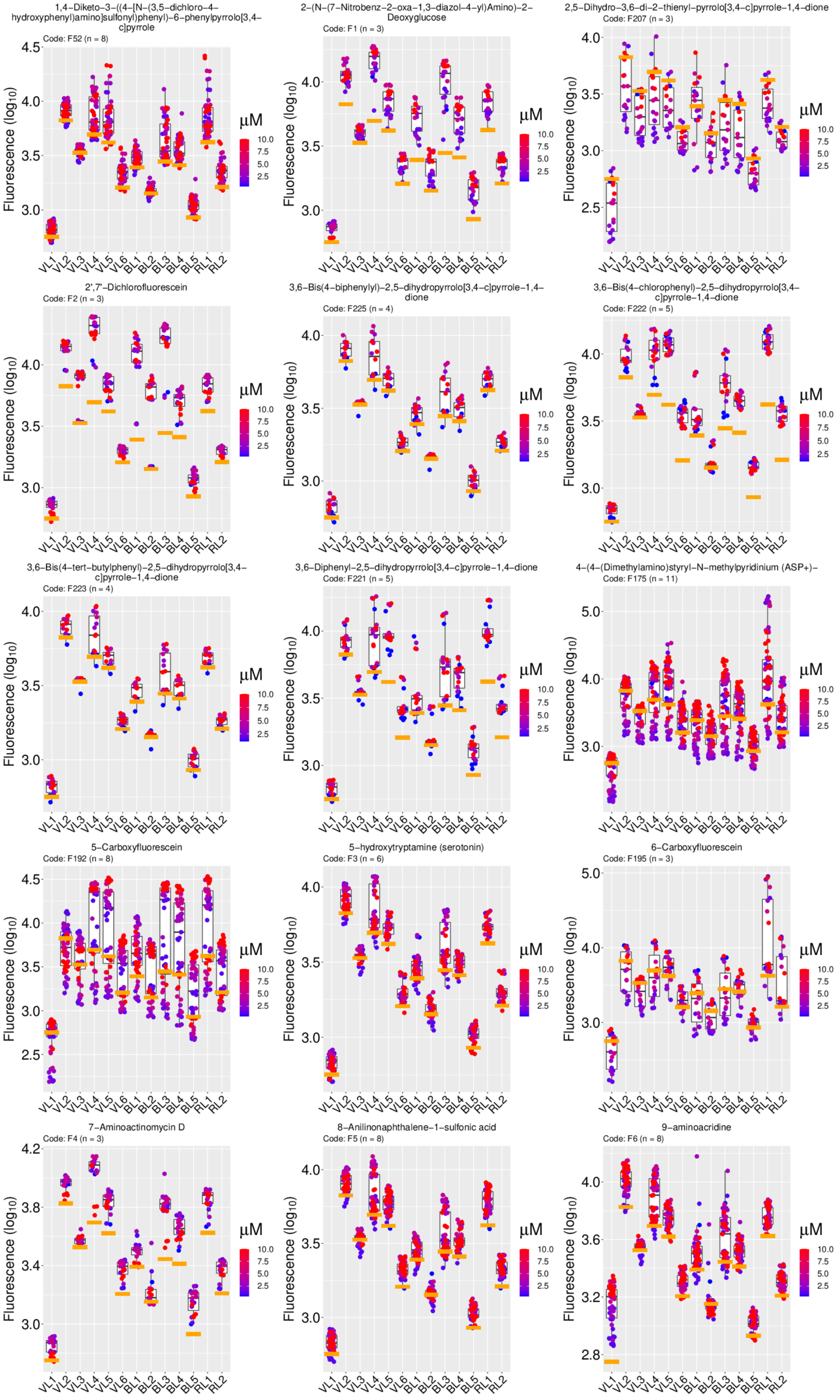

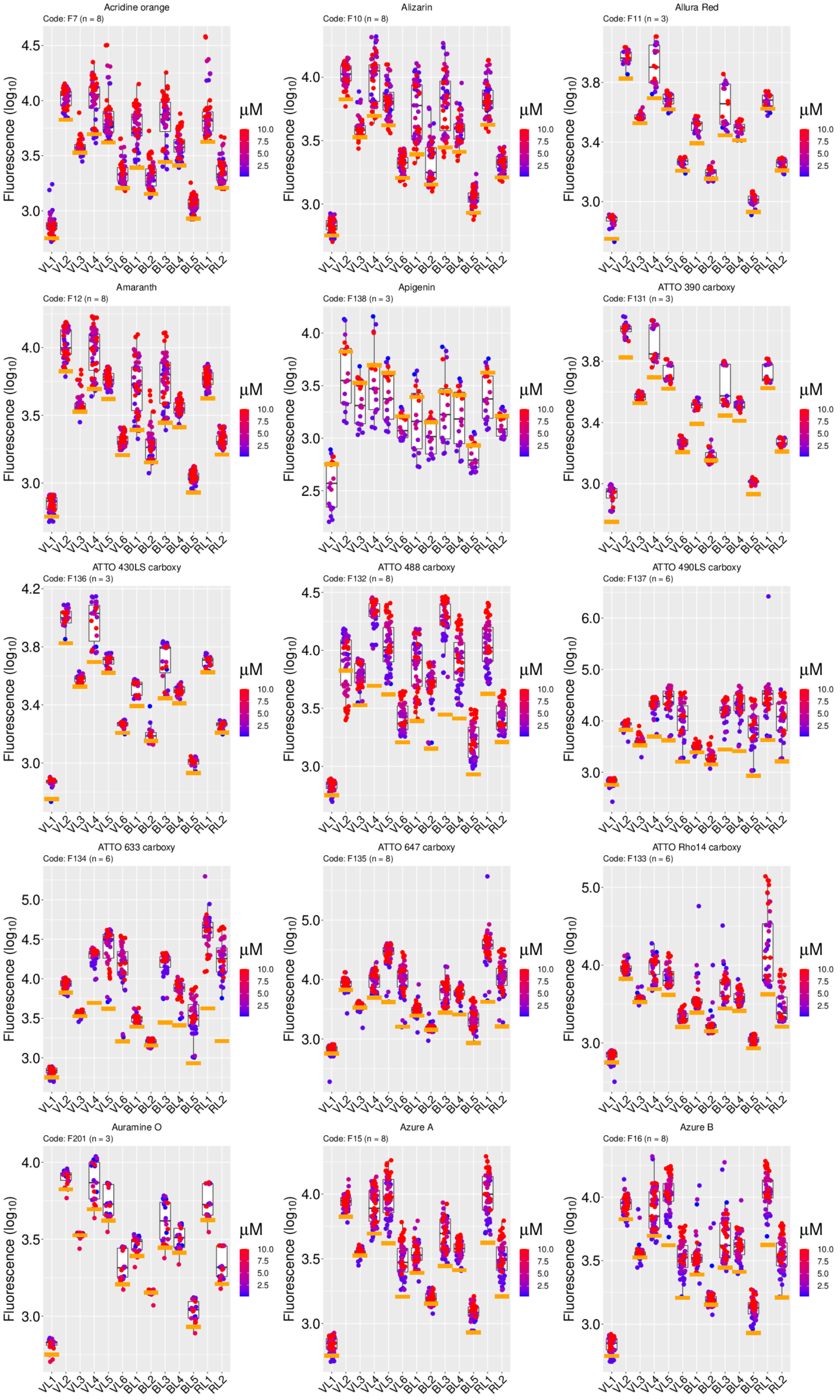

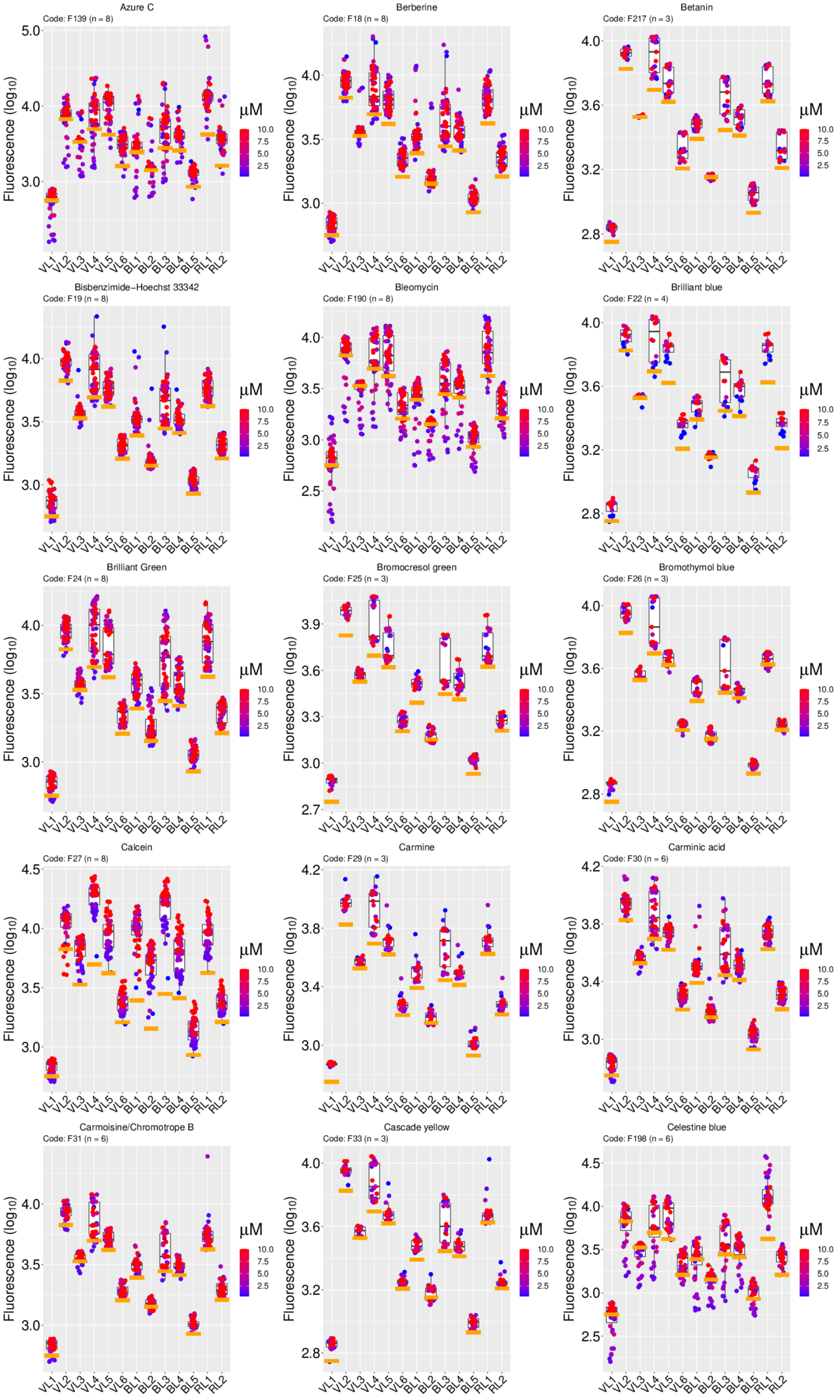

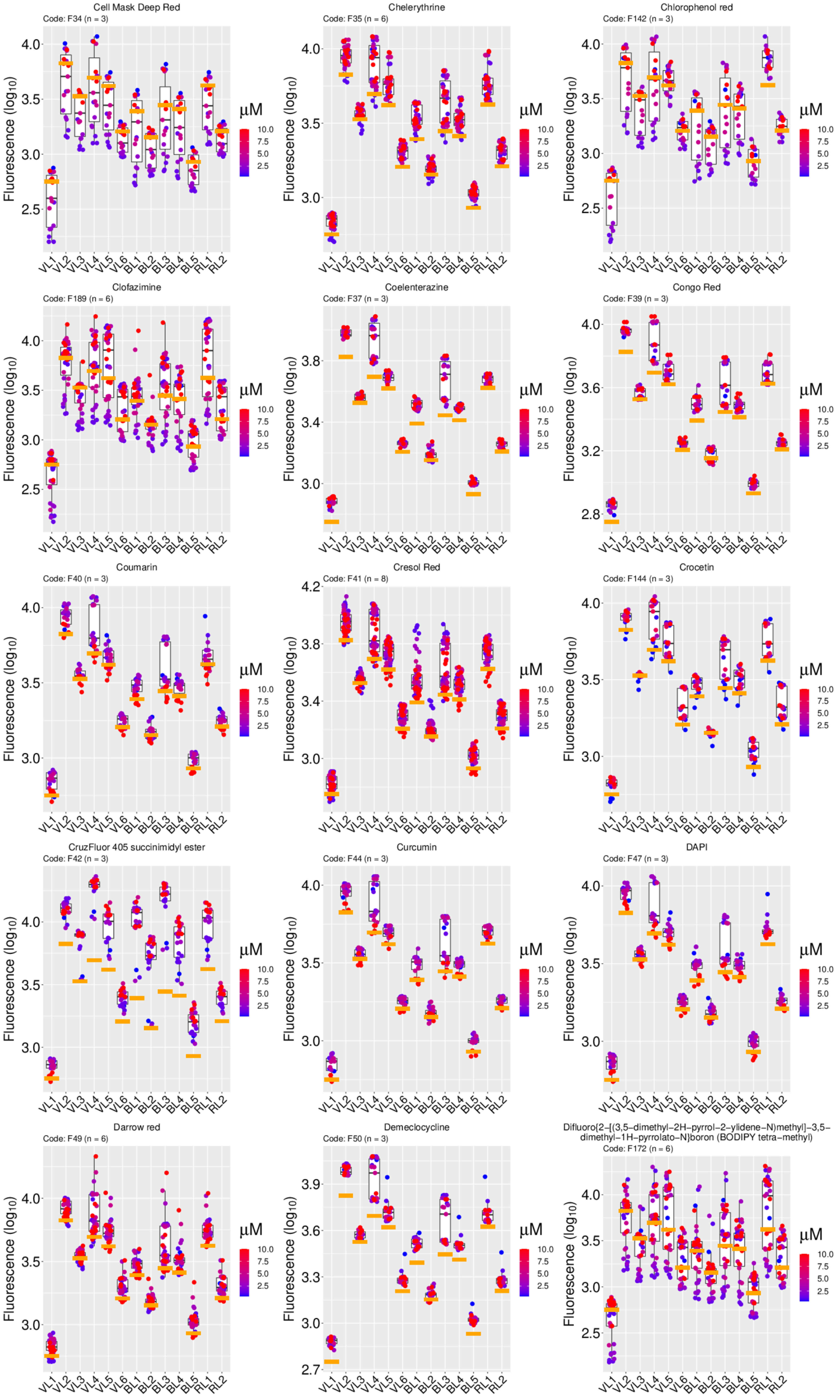

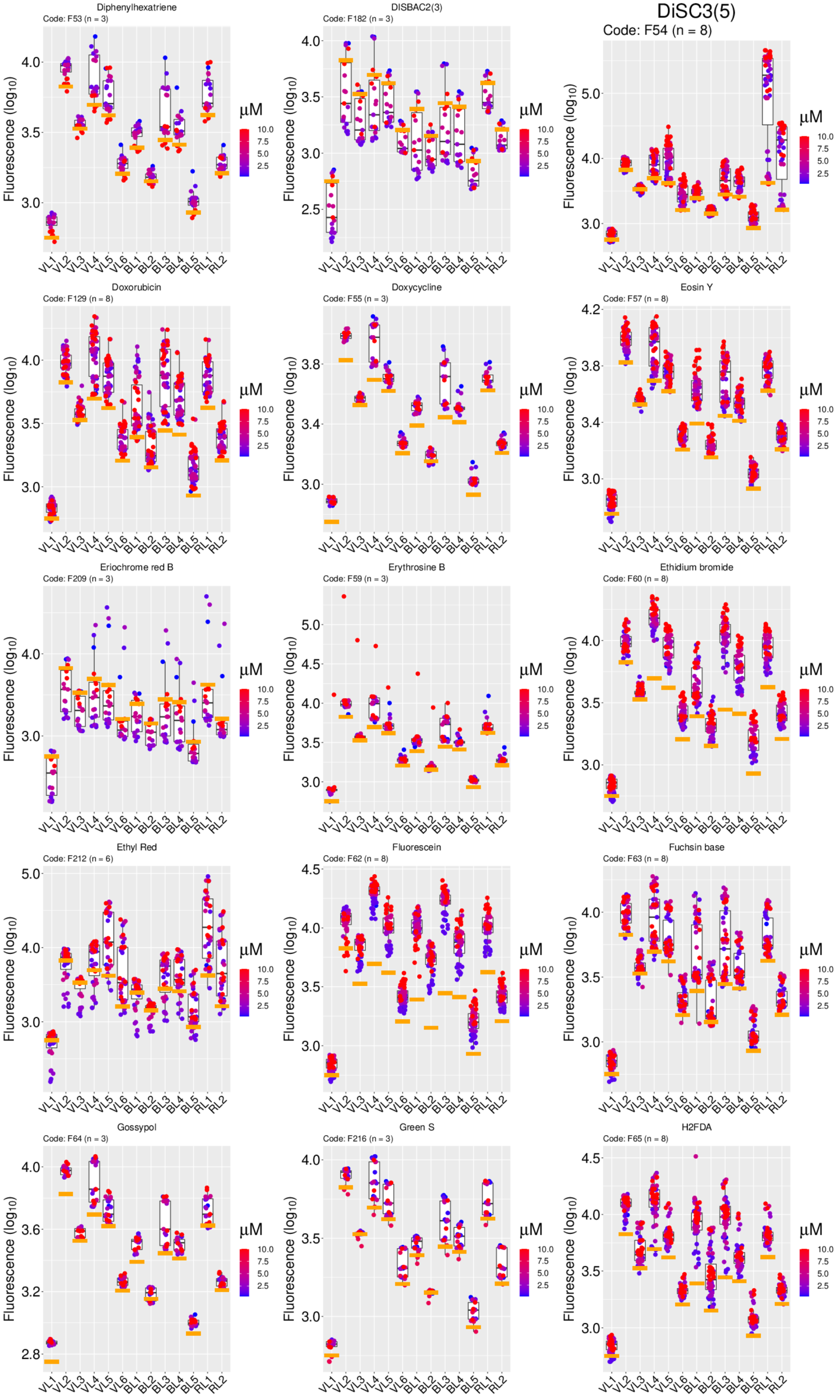

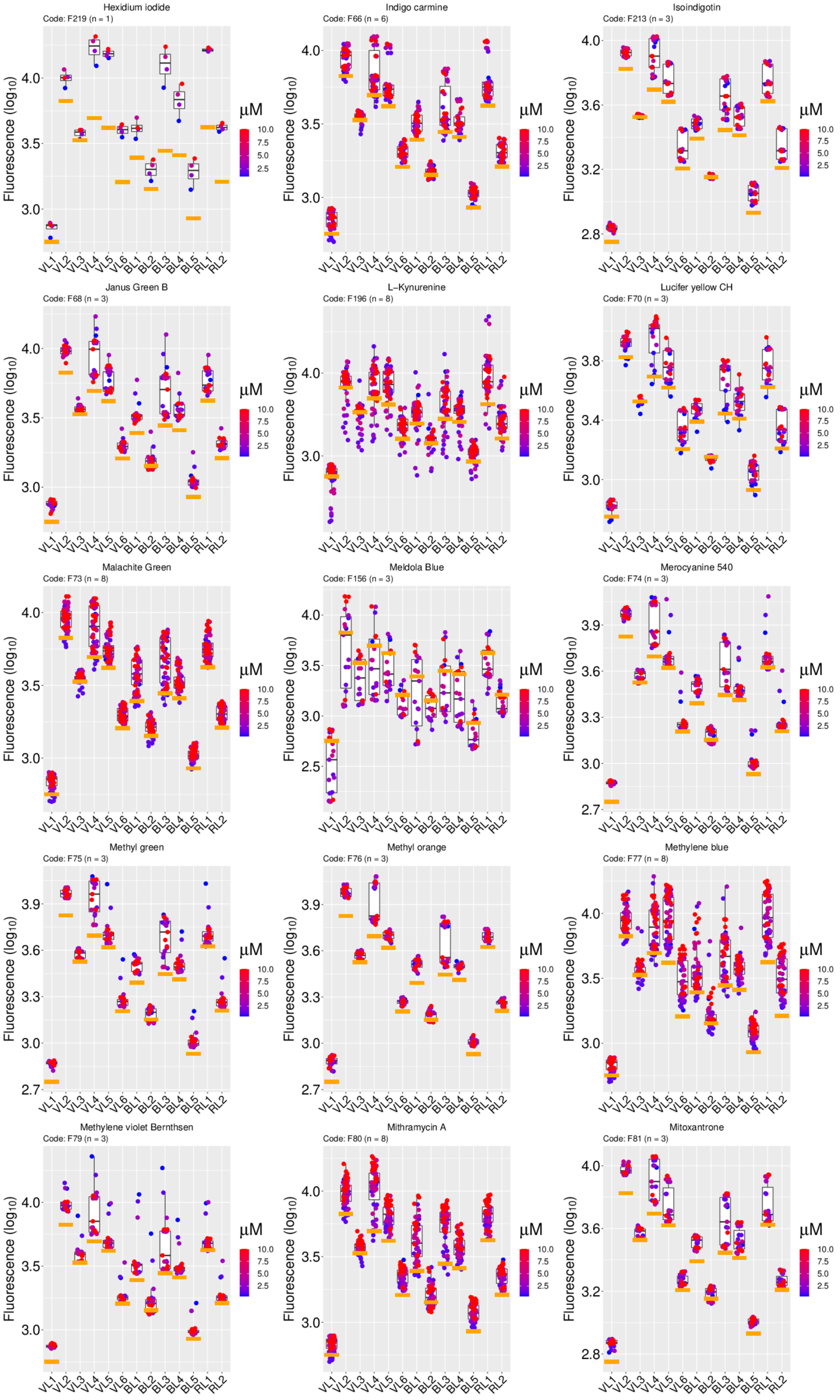

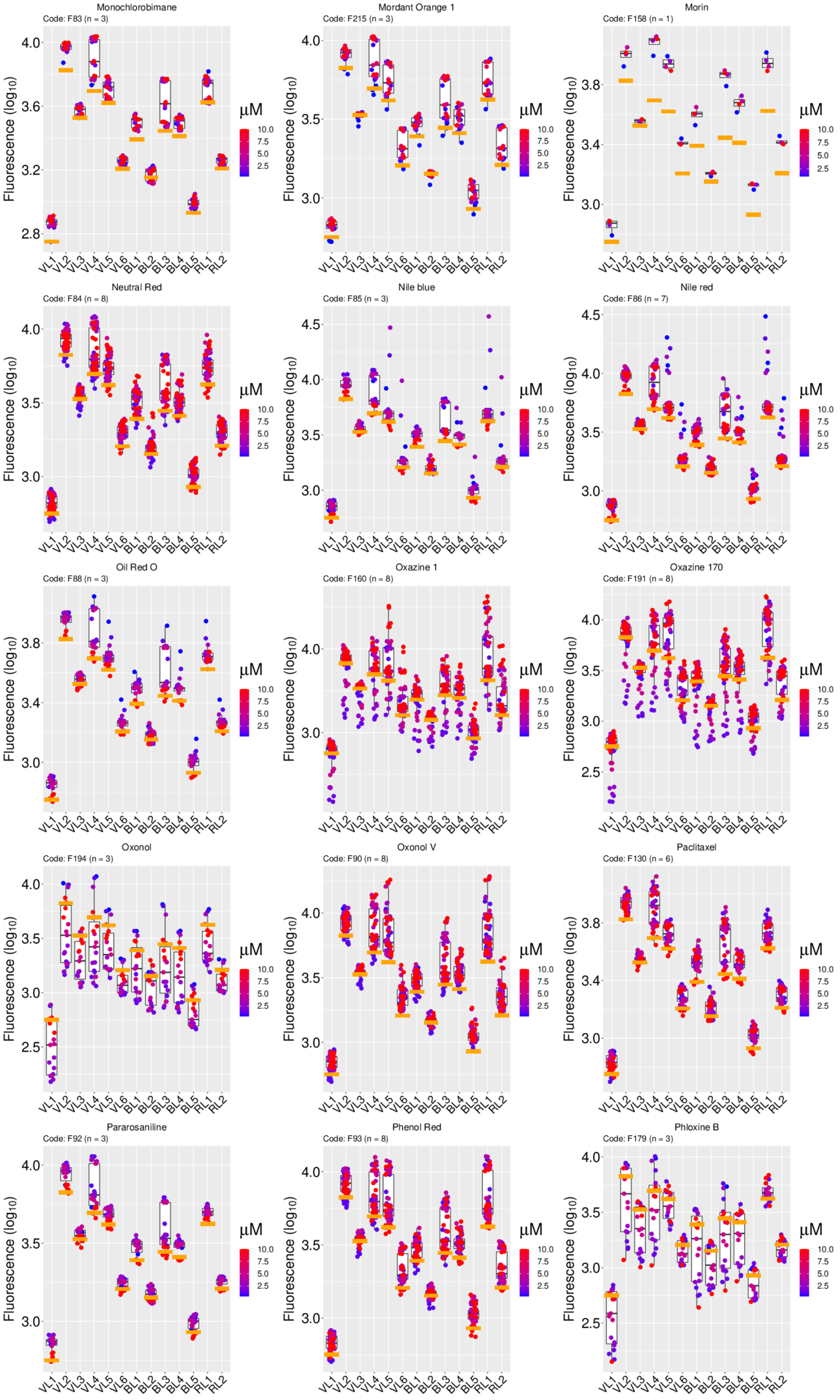

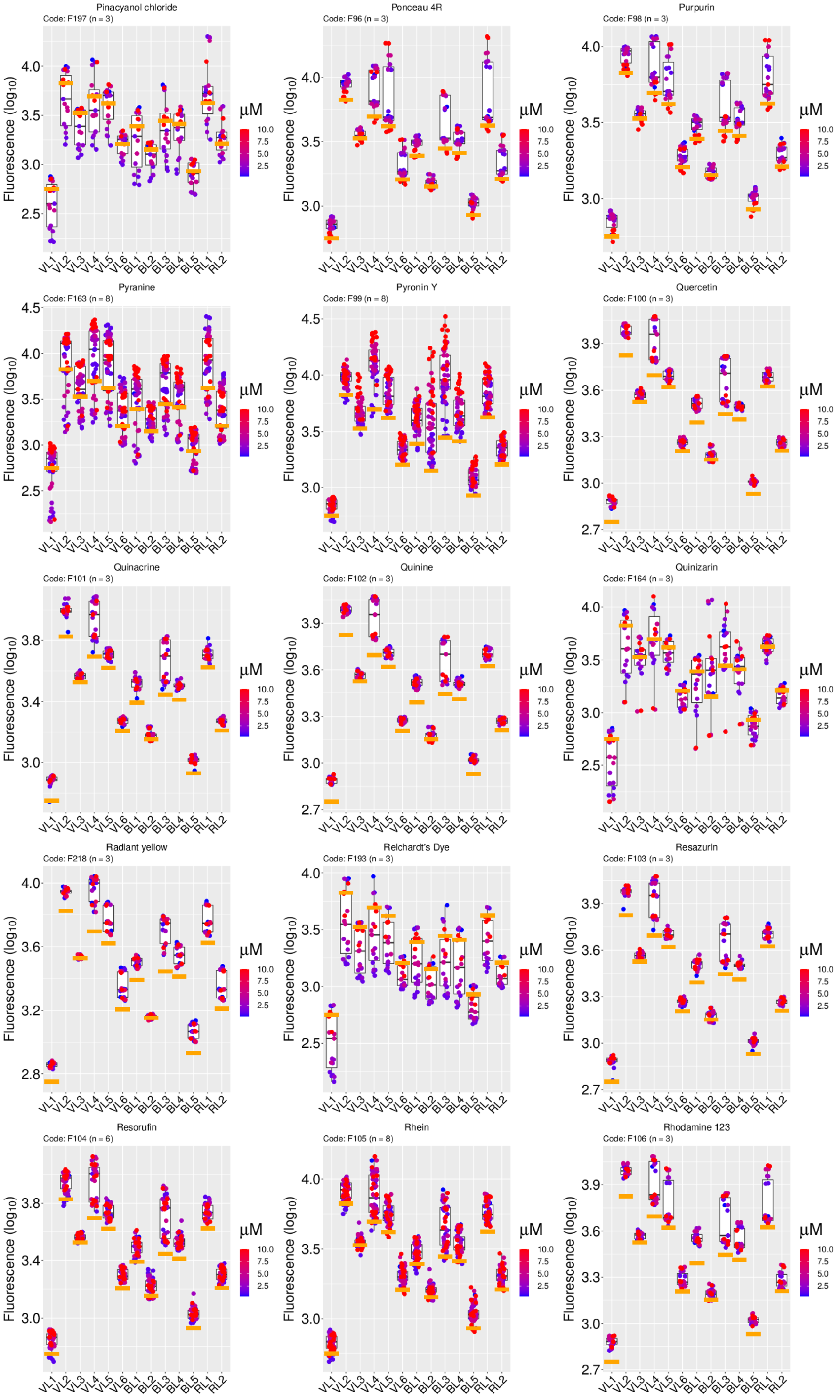

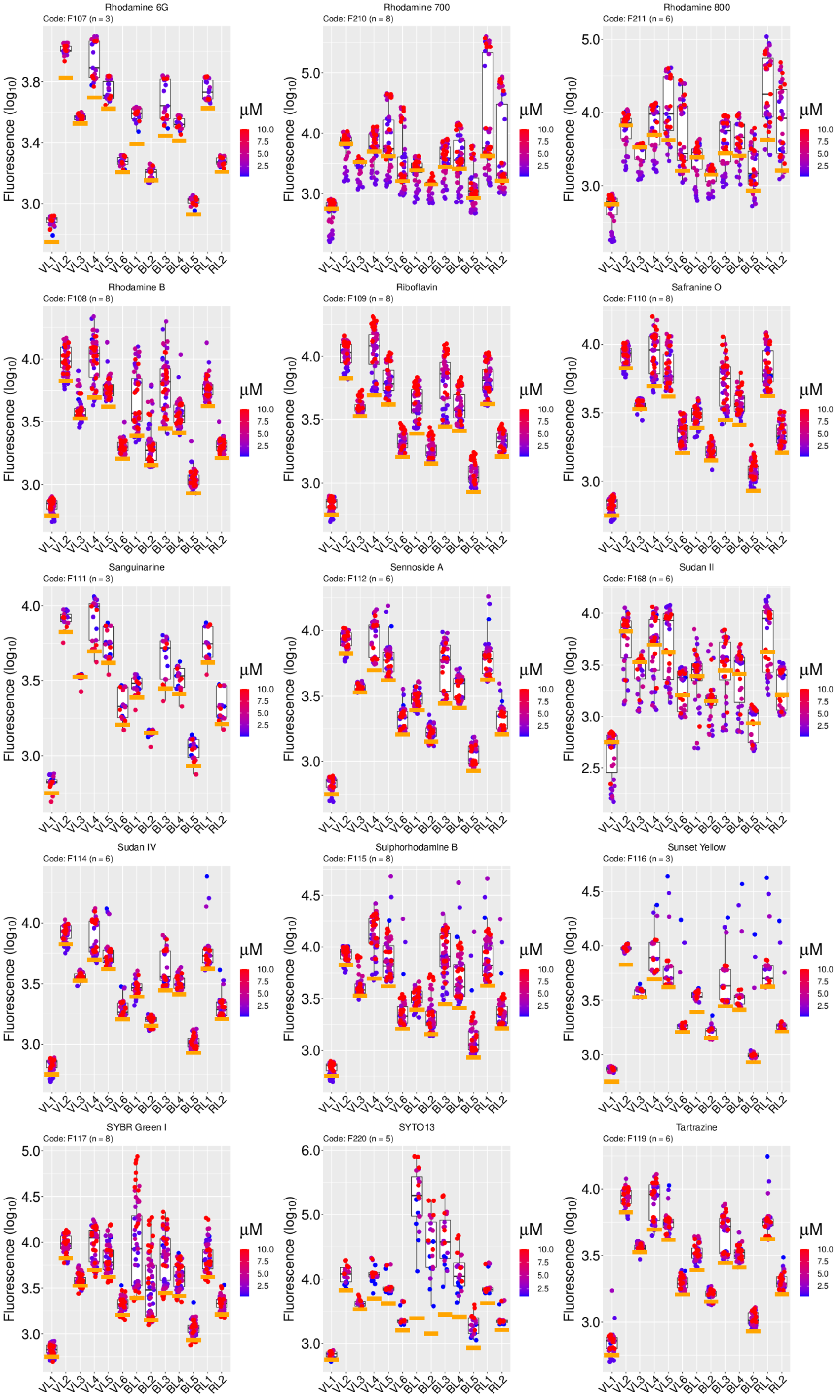

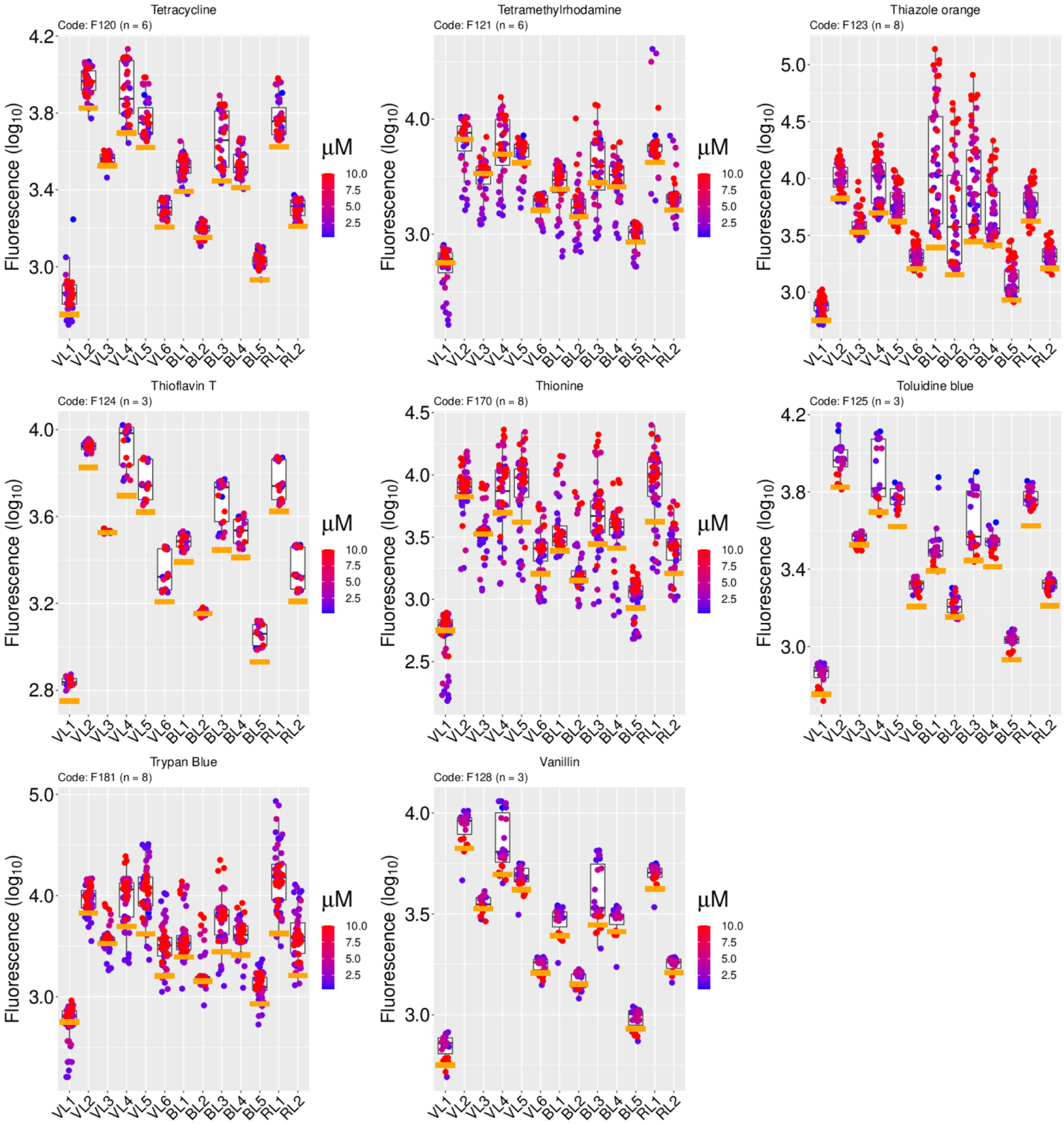

### Supplementary Fig 2

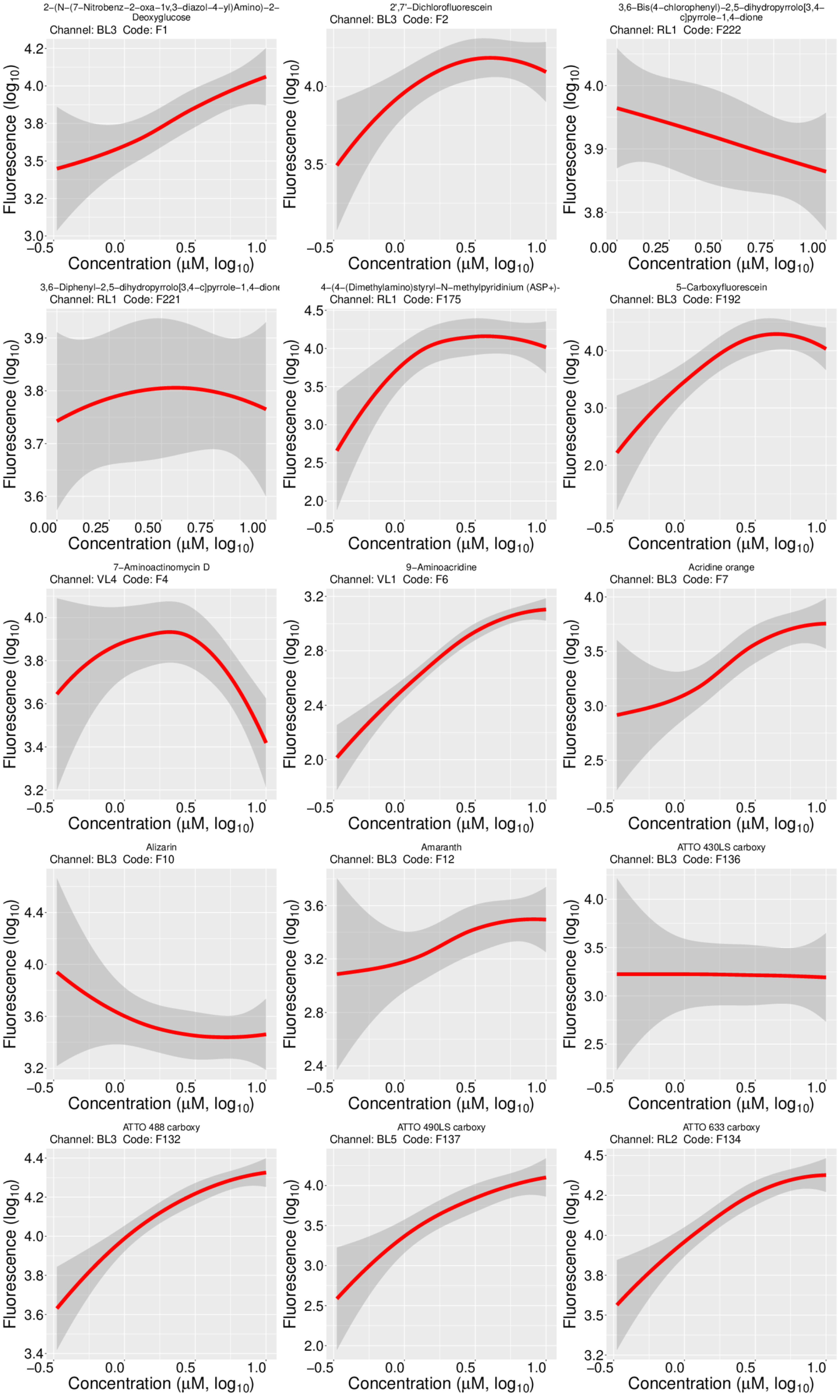

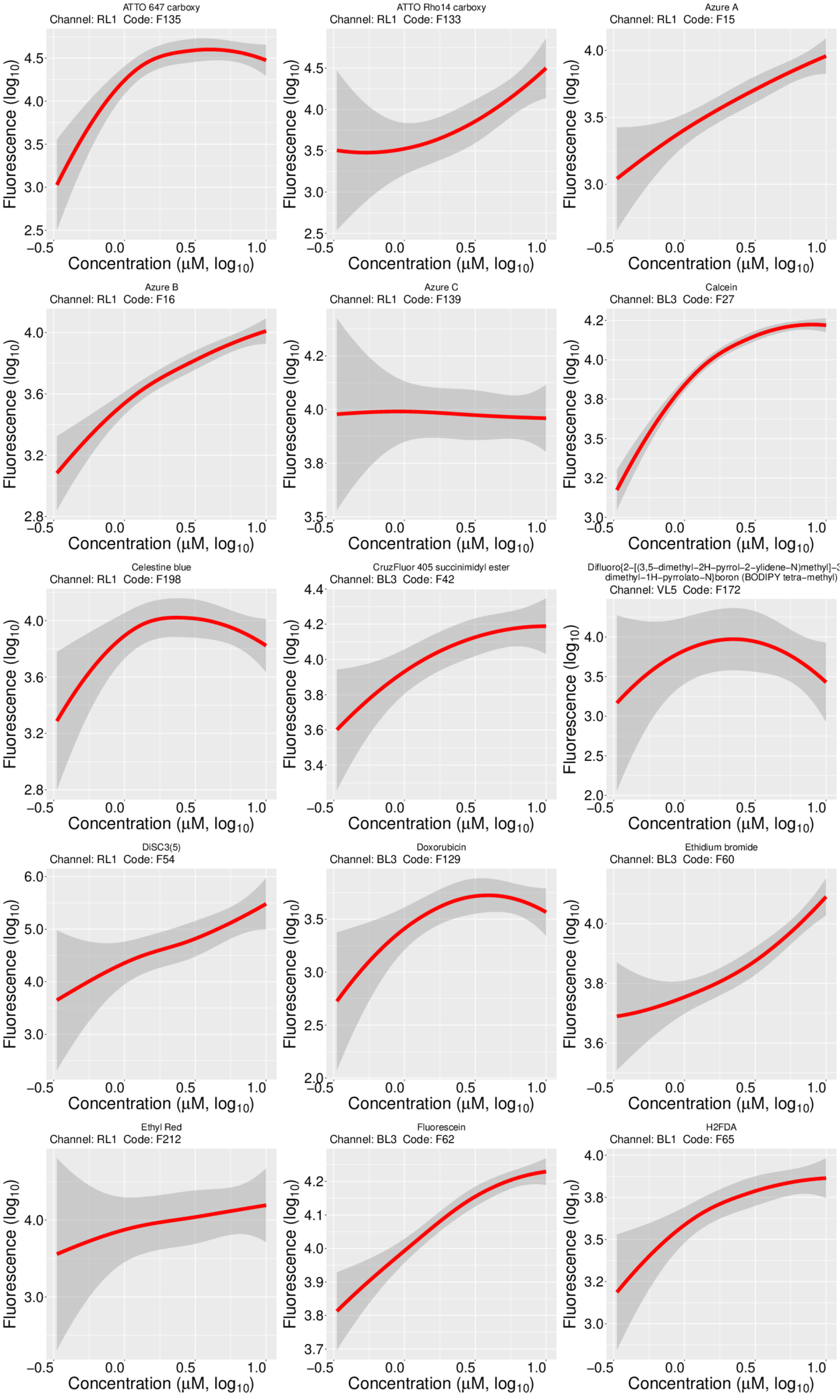

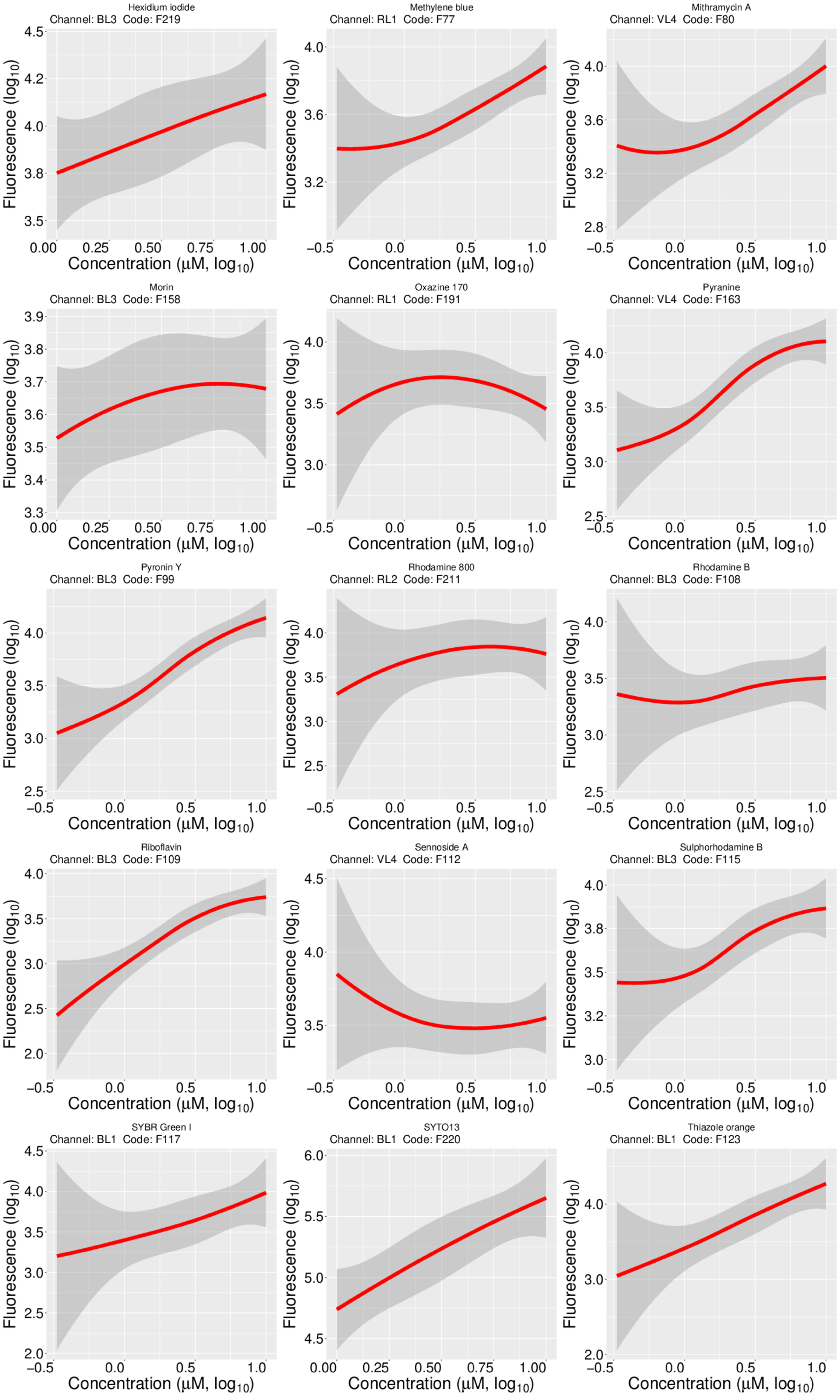

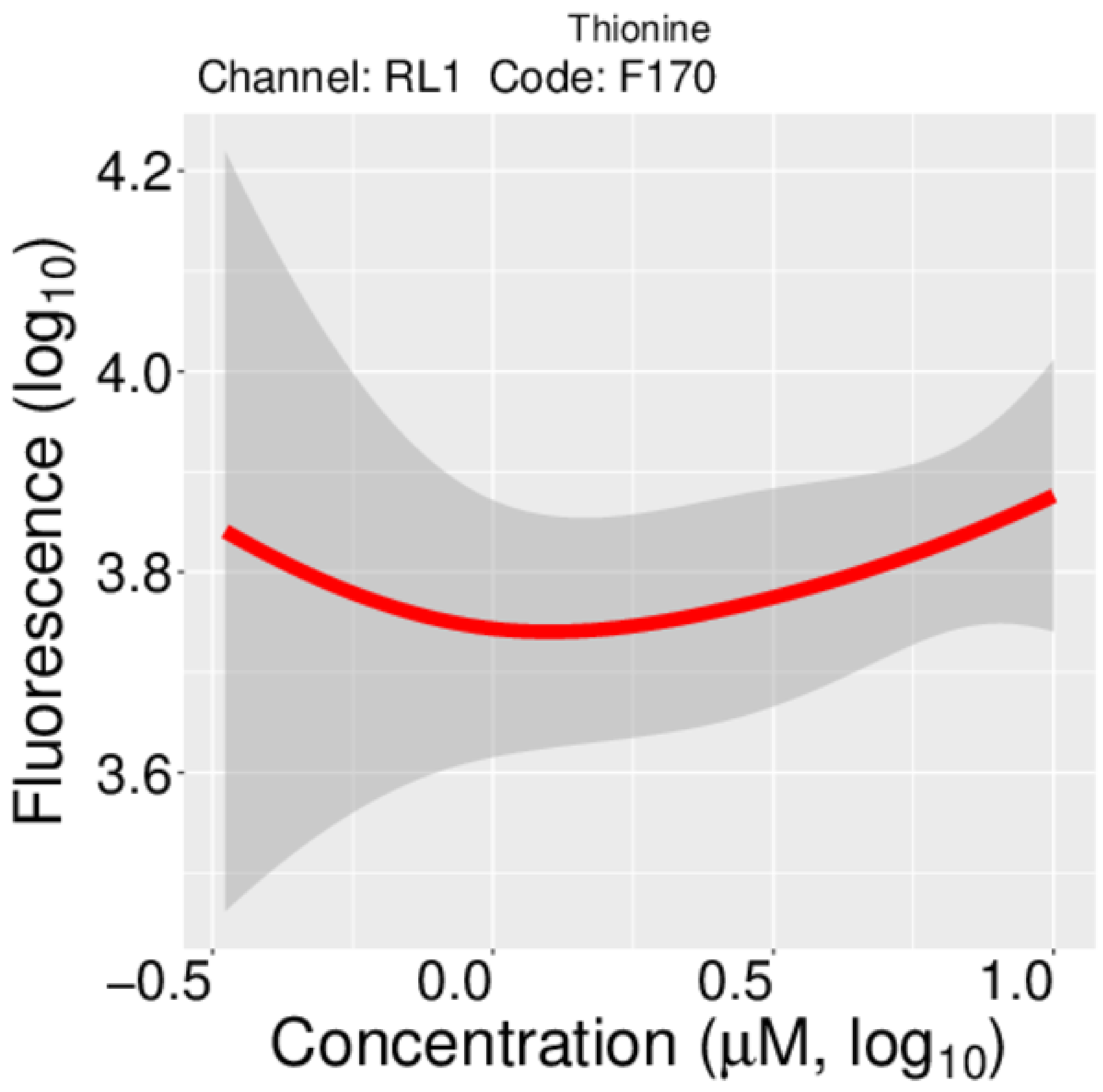

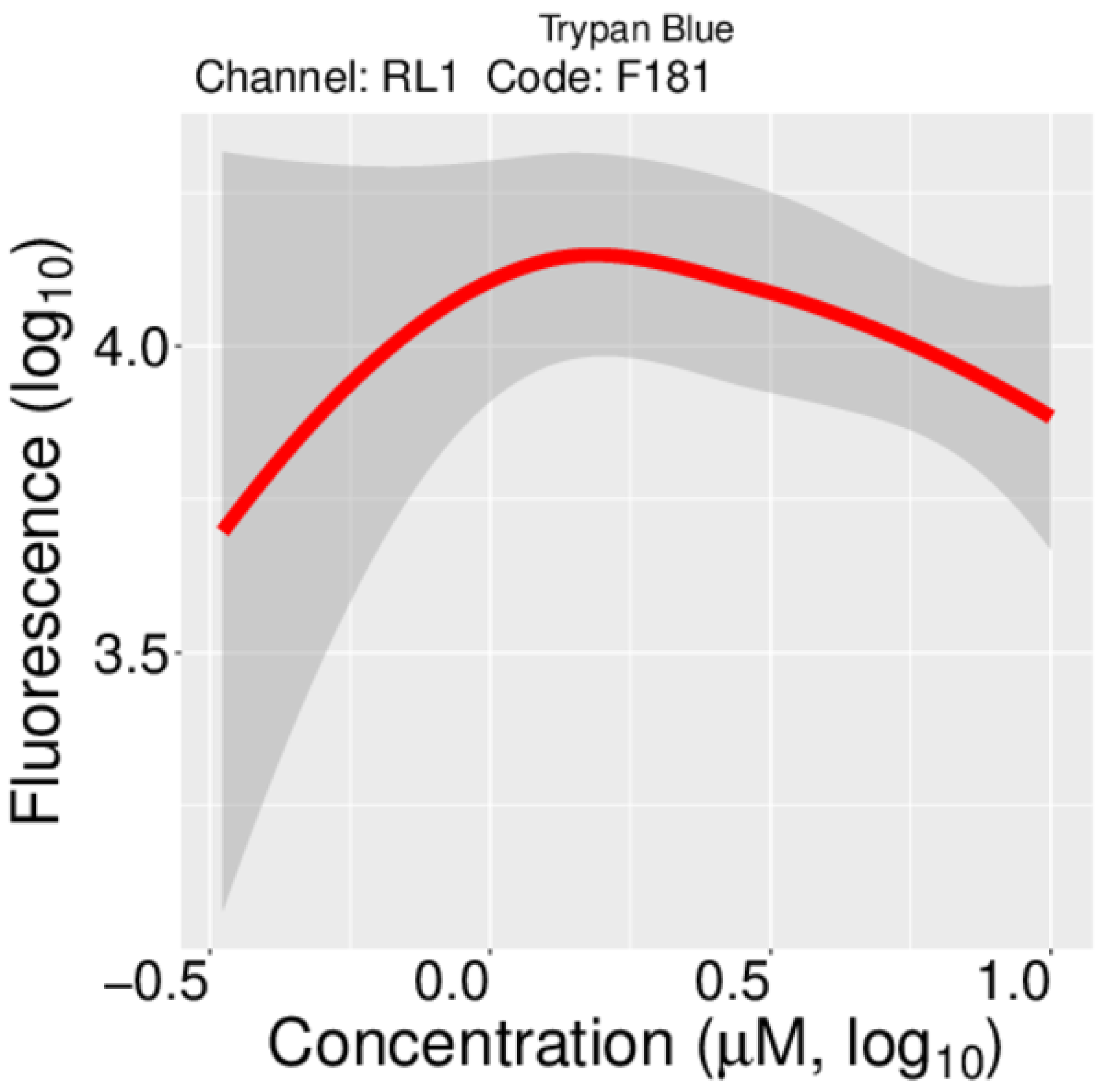

### Supplementary figure 3

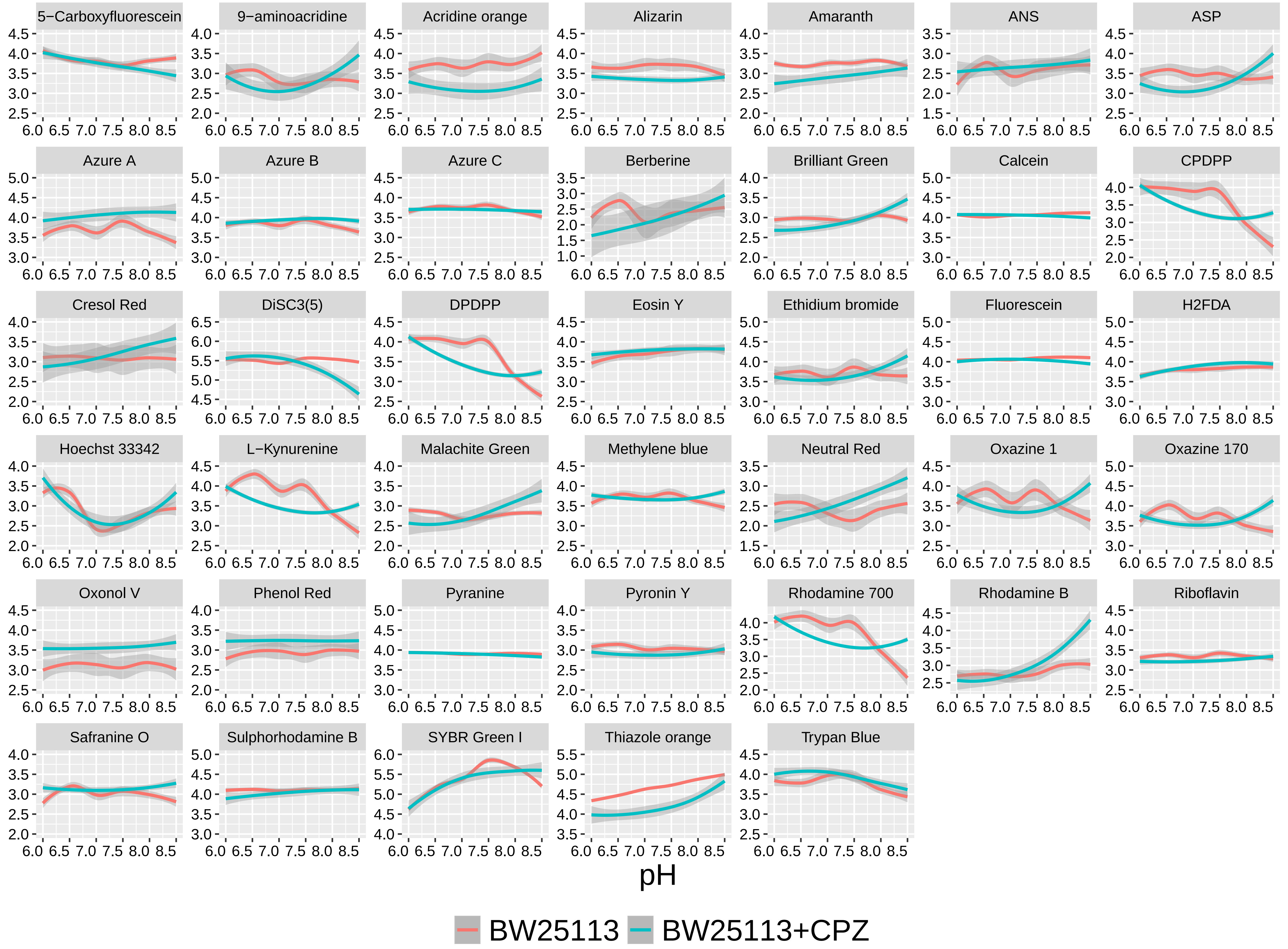
